# How Mycobacterium tuberculosis builds a home: Single-cell analysis reveals M. tuberculosis ESX-1-mediated accumulation of anti-inflammatory macrophages in infected mouse lungs

**DOI:** 10.1101/2024.04.20.590421

**Authors:** Weihao Zheng, Michael Borja, Leah Dorman, Jonathan Liu, Andy Zhou, Amanda Seng, Ritwicq Arjyal, Sara Sunshine, Alina Nalyvayko, Angela Pisco, Oren Rosenberg, Norma Neff, Beth Shoshana Zha

**Affiliations:** Division of Experimental Medicine, Department of Medicine, University of California, San Francisco, California, USA; Chan Zuckerberg Biohub, San Francisco, California, USA; Department of Biochemistry and Biophysics, University of California, San Francisco, California, USA; Division of Infectious Diseases, Department of Medicine, University of California, San Francisco, California, USA; Division of Pulmonary, Critical Care, Allergy and Sleep Medicine, Department of Medicine, University of California, San Francisco, California, USA

## Abstract

Mycobacterium tuberculosis (MTB) infects and replicates in lung mononuclear phagocytes (MNPs) with astounding ability to evade elimination. ESX-1, a type VII secretion system, acts as a virulence determinant that contributes to MTB’s ability to survive within MNPs, but its effect on MNP recruitment and/or differentiation remains unknown. Here, using single-cell RNA sequencing, we studied the role of ESX-1 in MNP heterogeneity and response in mice and murine bone marrow-derived macrophages (BMDM). We found that ESX-1 is required for MTB to recruit diverse MNP subsets with high MTB burden. Further, MTB induces an anti-inflammatory signature in MNPs and BMDM in an ESX-1 dependent manner. Similarly, spatial transcriptomics revealed an upregulation of anti-inflammatory signals in MTB lesions, where monocyte-derived macrophages concentrate near MTB-infected cells. Together, our findings suggest that MTB ESX-1 mediates the recruitment and differentiation of anti-inflammatory MNPs, which MTB can infect and manipulate for survival.

## Introduction

*Mycobacterium tuberculosis* (MTB) is the agent responsible for tuberculosis, one of the world’s deadliest infectious diseases.^1^ Central to the pathogenicity of MTB is its remarkable ability to survive and replicate within immune cells that are employed to destroy it. After inhalation, MTB enters alveolar macrophages (AMs),^2,3^ airway resident cells responsible for clearing inhaled pathogens and particles. In the early stages of infection, MTB-infected AMs adopt an anti-inflammatory signature that is less conducive to bacterial elimination.^2,4^ In this environment, MTB can replicate and induce the transfer of AMs to the interstitial compartment.^3^ Neutrophils and monocytes are recruited to the site of infection, and in parallel MTB-induced cell release induces transfer to newly recruited phagocytes, thus perpetuating its life cycle.^5–8^ An augmentation of data has begun to unveil the differential permissiveness of MTB survival in distinct subsets of mononuclear phagocytes (MNP) before and after the development of T cell responses, a potential key to the development of host-directed therapeutics to prevent MTB persistence in the lungs.^9,10^

Circulating blood monocytes, induced to egress from the bone marrow through CCR2 activation,^11^ predominantly exist as two distinct populations, Ly6c^hi^ (classical) and Ly6c^lo^ (non-classical), which correlate to CD14 and CD16 populations in humans.^12,13^ In the context of MTB-infected lungs in mice, Ly6c^hi^ monocytes represent the primary recruited population^14^ and differentiate into macrophages and dendritic cells (DC),^14–17^ which become the major infected cellular population.^18,19^ Control of infection can be accomplished after activation and recruitment of T cells, at which time bacterial burden begins to plateau but often does not result in eradication.^20–22^

Compelling evidence suggests that even after the development of T cell responses, specific subsets of recruited mononuclear phagocytes (MNP) are more permissive to MTB growth and replication than others.^3,18,19,23^ These findings have largely emerged by using a combination of flow cytometry and MTB expressing fluorescent proteins.^17^ However, flow cytometry’s limitations, such as the use of predefined antibodies, finite fluorophores,^24^ and variable gating strategies,^3,14,25^ have constrained the resolution of subset analysis and mechanistic understanding of survival of intracellular MTB. Furthermore, inherent challenges lie in reconciling cellular subsets between the MTB-infected environment, homeostasis, and other chronic inflammatory states due to cellular plasticity and heterogeneity. For instance, during MTB infection, AMs can upregulate MHCII.^19^ In addition, CD11b^+^CD11c^+^ cells, previously defined as monocyte-derived DC,^14,17,26^ are heterogeneous and include monocyte-derived macrophages.^19^

The application of single-cell RNA sequencing (scRNA-seq) has provided a wealth of information that is instrumental in unraveling the complexity of cell heterogeneity in unbiased standards, illuminating organ composition,^27,28^ characterization of unique cell types,^29,30^ and delineating host cellular changes in the presence of an infecting organism.^31^ With this technology, there is increasing evidence of the complexity of MNPs in the lungs, leading to refined definitions of recruited and resident interstitial macrophages (IM) and DC populations.^32-35^ Understanding subsets of recruited MNPs holds the promise of elucidating cell types that differ in their permissiveness for MTB, allowing for a mechanistic understanding of how MTB can sustain itself in the cell and holds potential for targets of host-directed therapies.

In addition to surviving in MNP, MTB employs other mechanisms to persist and cause disease, including cell-to-cell transfer,^7^ induction of cellular recruitment,^36^ and delaying activation of the adaptive immune response,^26^ collectively conferring MTB with a sustainable niche. One central virulence factor of MTB that is involved in key components of pathogenesis is early secretory antigenic target secretion system-1 (ESX-1), a type VII secretion system^37^ shown to be involved in multiple pathways including phagosome escape,^38,39^ inhibition of phagolysosome maturation,^40^ and recruitment of macrophages.^41,42^ Components of the ESX-1 machinery are primarily encoded in the operon region of difference 1 (RD1), which is lacking in the vaccine strain, bacille Calmette-Guerin (BCG).^43^ While attenuated, MTB lacking functional ESX-1 still survive, replicate, and spread cell-to-cell in mice after aerosol infection,^37,44^ but induce less myeloid cell recruitment.^7,18^ We hypothesize that ESX-1 drives recruitment of MNP since it is vital to pathogenesis but not essential for the organism’s survival.

Here, we aimed to elucidate the role of MTB ESX-1 in MNP recruitment and heterogeneity, identify permissive subsets of MNP, and understand the spatial interaction of recruited bystander MNP, infected MNP, and the bacteria. Using aerosol infection, we compared uninfected mice, mice infected with MTB H37Rv, and mice infected with an attenuated H37Rv lacking functional ESX-1 machinery. We find that ESX-1 deficient H37Rv does not recruit and induce activation of macrophages to the same extent as the wild-type strain. Using both in vitro and in vivo models, we found that ESX-1 is important in driving an anti-inflammatory response to manipulate a host cell for MTB’s survival. Using spatial transcriptomics, we also found an upregulation of anti-inflammatory signals in MTB lesions, where monocyte-derived macrophages concentrate near MTB-infected cells.

## Results

### Mycobacterium tuberculosis recruits heterogeneous subsets of mononuclear phagocytes

Previous work has demonstrated that, following MTB infection, monocyte-derived (moDerived) cells are recruited to the lungs and undergo differentiation over time.^14,25^ Recent work demonstrates that soon after the development of T cell responses, monocyte derived lung cells (MNC) are more permissive to MTB survival compared with AM.^18,19^ Despite best efforts at refined profiling, these cell subsets defined by flow-cytometry remain heterogeneous. To investigate the differential properties of these and other MNP cells in the early chronic infection,^45^ we utilized single-cell RNA sequencing. Mice were infected with H37Rv expressing mCherry and euthanized 28 days later to allow development of the adaptive T cell response. After gating out NK, T, and B cells, live cells were sorted by mCherry expression (i.e. positive indicative as infected or negative indicative as bystander; SuppFig1A). Cells were processed using the 10X genomic platform on live cells within a BSL3 (Fig1A). Hierarchical Leiden clustering of cellular transcriptional libraries revealed multiple discrete cellular populations, however, heterogeneity of cell subsets was not clearly delineated despite computational sub-clustering. We then used SmartSeq2, a plate-based scRNA-seq platform in which single cells are directly sorted into gentle lysis buffer.^46^ To capture cells of highest interest, single lung cells were sorted for CD11b and/or CD11c positivity into 384-well plates based on mCherry expression (SuppFig1B, Fig1A). As compared to 10X, SmartSeq2 demonstrated higher total counts with less ribosomal RNA capture as well as a clearer delineation of unique moDerived cell subtypes (SuppFig2).

**Figure 1:**
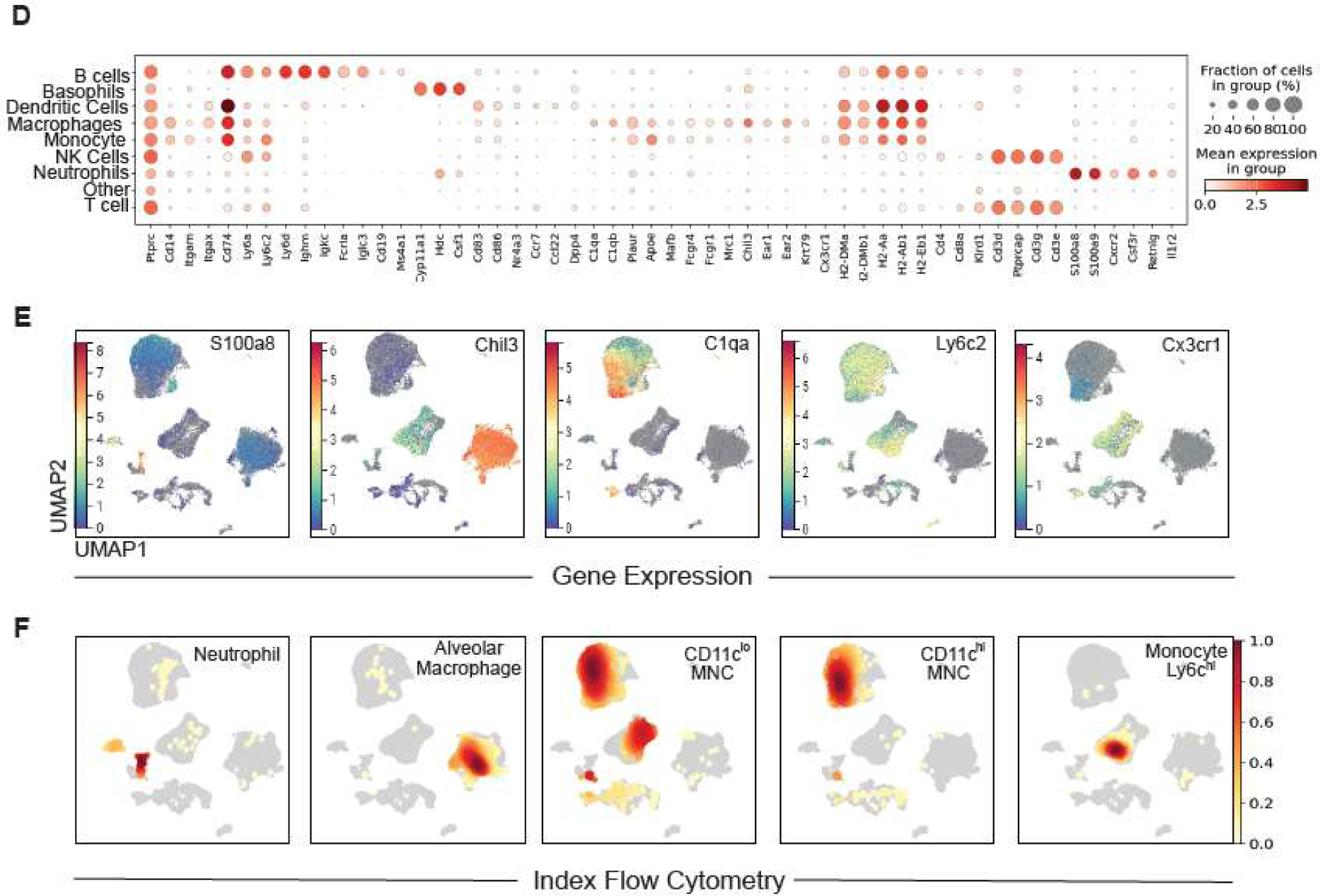
Mycobacterium tuberculosis induces recruitment of heterogeneous MNPs in the lung of mice. **A)** Mice were infected with H37Rv for 28 days, lung dissociated to single cell suspension, and live cells flow sorted to obtain myeloid cells differentiated by infection status based on bacterial fluorescence expression (SuppFig 1). Cells were either sorted directly into 384 well plates (1; SS2: SmartSeq2) or washed and counted for 10X chemistry (2). After reverse transcription and decontamination, samples were removed from the BSL3 for library preparation and sequencing. **B)** Ingest was used to combine SmartSeq2 and 10X sequencing datasets into single UMAP plot. **(C)** Cellular subsets were classified using the Immgen database within SinglR for unbiased annotation. **(D)** Dot plot shows transcriptional canonical markers where fraction of cells expressing the gene is denoted by size of dot and the darker red denotes higher mean gene expression of the gene. **(E)** Gene expression of key canonical markers after log transformation and normalization visualized using CellxGene, where darker red signifies higher expression. **(F)** Density plots of subsets obtained through index sorting for SmartSeq2 using the mean fluorescent intensity of surface markers were converted using boolenization and overlayed onto the transcriptional data.

Given the higher number of cells captured, sequencing data obtained from SmartSeq2 were integrated with 10X for unbiased annotation using SinglR with mouse ImmGen database as reference^29^ (Fig1B,C). Nomenclature was confirmed by expression of key canonical transcriptional markers visualized in sequence (Fig1D) and with use of overlay on UMAP by Cellxgene^47^ (Fig1E). In addition, index sorting during SmartSeq2 provided single cell mean fluorescent intensity (MFI) analysis of antigen abundance. Mean fluorescence intensity (MFI) of key surface markers used during gating was overlayed onto transcriptional cell clusters using Boolean computing (Fig1F, SuppFig1D). Clear identification of alveolar macrophages (CD11c^+^SiglecF^+^, *Chil3*), neutrophils (Ly6g^+^, *S100a8* high), and monocytes (Ly6c^hi^, *Cx3cr1*+) could be delineated. Further, the majority of CD11c^hi^-MNC corresponded to the transcriptional annotation of macrophages, while CD11c^lo^-MNC contained both macrophages and monocyte-like cells. Together, index sorting and SmartSeq2 confirm the identification of heterogenous MNP at both mRNA and protein levels in early chronic MTB-infected mouse lungs.

### Lack of a functional ESX-1 prevents recruitment of diverse monocyte-derived macrophages

ESX-1 plays an important role in myeloid cell recruitment and bacterial cell-to-cell transfer. However, mice can successfully be infected with MTB lacking functional ESX-1.^7,18,37^ We next asked if functional depletion of ESX-1 would result in alteration of the observed heterogeneous recruitment of MNP, which may be a cause of alteration of the dynamics of bacterial cell-to-cell transfer.^7,18^ *EccD*_1_, a critical component of the ESX-1 complex formation,^48^ was deleted using phage-mediated allelic exchange,^49^ and bacteria were confirmed to no longer secrete CFP-10 and ESAT-6, despite expression of the proteins (SuppFig3A,B). When used to infect mice, MTB lacking EccD_1_ resulted in fewer CD11c^hi^-MNC, CD11c^lo^-MNC, and neutrophils by flow-cytometry (Fig2A,B), comparable to findings when region of deletion 1 (RD1) is deleted.^18,50,51^

**Figure 2.**
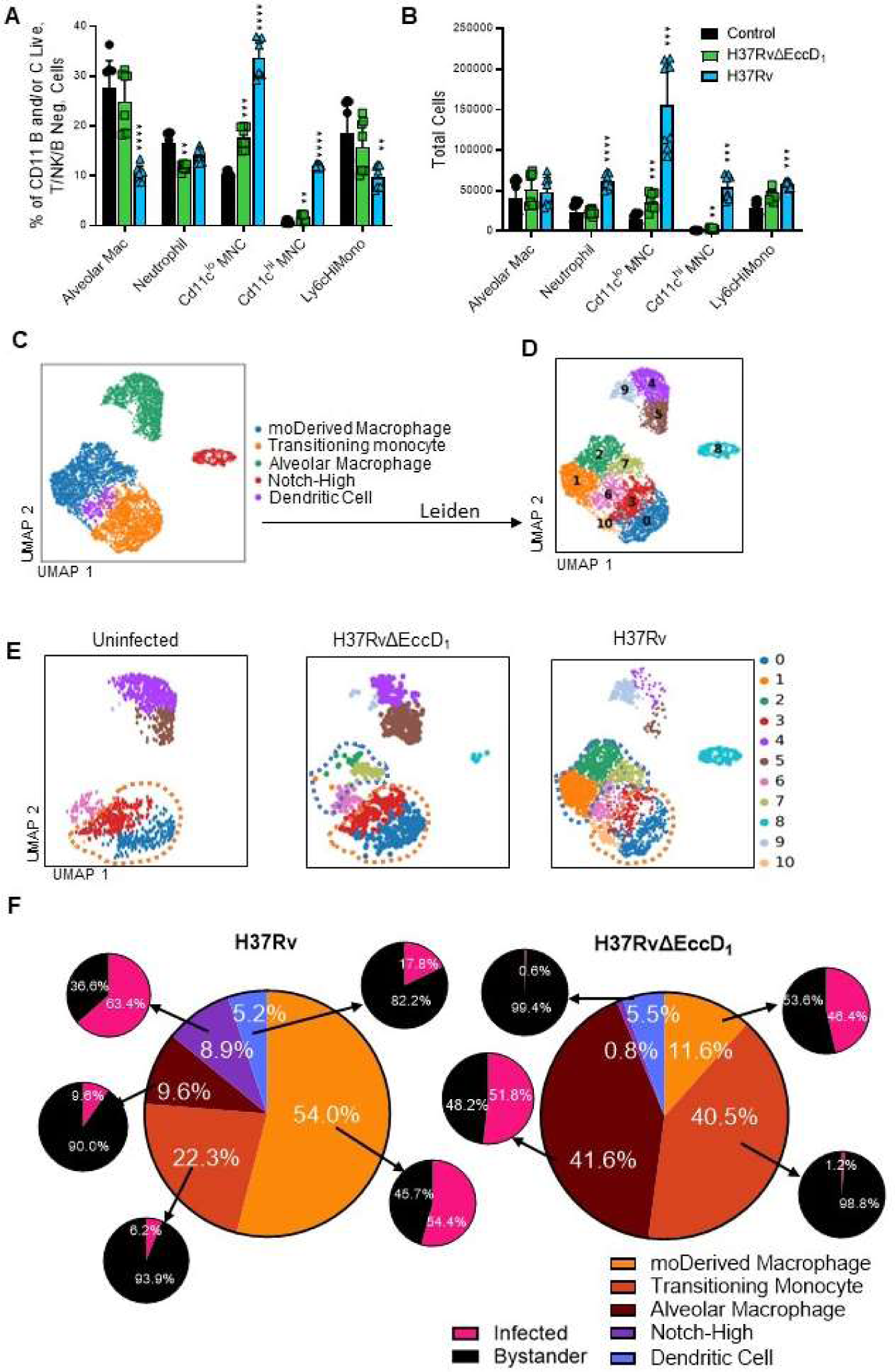
SmartSeq2 reveals ESX-1 recruitment of diverse subsets of monocyte-derived macrophages. **A,B)** C57BL/6 mice were infected for 28 days with H37Rv or H37RvΔEccD_1_ and lung mononuclear phagocytes were analyzed by flow cytometry compared to uninfected control mice. **C)** SmartSeq2 analysis of CD11b and/or CD11c positive cells from mice infected with H37Rv or H37RvΔEccD_1_. **D)** Cellular subsets were further clustered using Leiden algorithm. **E)** Separation by infection status of the mouse that the cells were isolated, with moDerived macrophages denoted by blue hashtag and transitional cells by an orange hashtag. **F)** Percent of subsets recruited were graphed by parts of a whole per infected strain. Percent of cells infected was calculated per cellular subset and denoted by the subset graph of pink (infected) and black (bystander).*p<0.05, **p<0.01, ***p<0.001, ****p<0.0001 using t-test with Welch correction and Holm-Sidak multiple comparisons.

Given that cells infected with H37Rv lacking the operon RD1 during aerosol infection are rarely sorted,^7^ we employed SmartSeq2 to capture rare events and thoroughly analyze recruited cell subsets within the heterogenous MNC population. Mice were infected with MTB expressing zsGreen (SuppFig1C). To ensure aligned cellular identification and analysis, transcriptomic data was overlayed using Ingest (Scanpy) with our atlas (SuppFig3C), with cell subsets confirmed as identified above. In line with flow cytometric results, the proportion of cell types present depended on infection status, with the majority of cells identified as alveolar macrophages in the uninfected mouse, recruited MNPs from H37Rv-infected mice, and a mixture in the proportion of alveolar macrophages and recruited cells in H37Rv ΔEccD_1_-infected mice.

Monocytes, macrophages, and dendritic cells from both SmartSeq2 experiments were subset from the larger atlas to standardize method, sequencing depth, and further factors that cannot be compensated computationally (SuppFig3E). Cells were reclustered and noted to have similar distributions as the combined cell atlas. With this method, 5 clusters were identified: moDerived Macrophages, transitioning monocytes, alveolar macrophages, moDerived cells with elevated Notch signaling and underrepresentation of canonical markers (‘Notch-High’), and a smaller DC population. Nomenclature was again confirmed by transcriptional canonical markers and previously defined surface markers^18^ (Fig2C, SuppFig4A-C). For instance, the denotation transitioning monocyte was given due to a high expression of monocyte markers (e.g. *Cx3cr1*, *Sell, Spn, Plac8, Nr4a1*) with further signature of *C1q*, *Cd68*, *Csf1r*, *Lyz2* (SuppFig4B). Given experimental methods to exclude intravascular monocytes combined with transcriptional similarities, those with hyper Ly6c surface expression are also expected to be recently recruited or patrolling monocytes.^52^ Conversely, moDerived macrophages demonstrated significantly lower expression of monocyte markers while retaining *Ly6c2, Cd14*, MHC antigen presentation, and *C1q* markers.

Further subsetting was accomplished through Cellxgene and Leiden^53^ clustering (Fig2D), revealing 3 subsets each of moDerived macrophages (cluster 1, 2, 7), transitioning monocytes (cluster 0, 3, and 10), and AMs (cluster 4, 5, 9), each with differential infection states. As hypothesized, mice infected with H37RvΔEccD_1_ had significantly fewer infected cells compared to H37Rv, with the majority of infected cells clustered within AMs (Fig2E,F). In contrast, the major infected subset is moDerived macrophages from the lungs of H37Rv-infected mice, consistent with a previous study.^18^ Notably, there were significantly fewer moDerived macrophages in the lungs of H37RvΔEccD_1_ infected mice (SuppTable1). Similarly, there was a reverse in proportion of moDerived macrophages to transitioning monocytes in H37RvΔEccD_1_-infected mice (40.5% versus 11.6% of all cells, p<0.0001) as compared to H37Rv-infected (22.3% versus 54% of all cells, p<0.0001). Therefore, lack of ESX-1 resulted in a lack of moDerived macrophages despite recruitment of monocytes.

### H37Rv with functional ESX-1 induces recruitment of anti-inflammatory macrophages with high MTB burden

We next turned to understand the transcriptional differences of MNPs recruited during MTB infection. Differentially expressed genes between cluster and subclusters were determined using MAST (R), with gene ontology and pathway enrichment analysis conducted with genes meeting a false-discovery rate of <0.05 (SuppTable2). Supporting identified cell types, Qiagen Ingenuity Pathway Analysis (IPA) revealed signatures of oxidative phosphorylation, necroptosis, and autophagy in the moDerived macrophage cell cluster compared with transitioning monocytes. Conversely, transitioning monocytes were enriched for pathways important in translation (i.e. EIF2) and cellular movement (i.e. integrin signaling and extravasation) (SuppFig3E).

Gene ontology analysis was conducted between cell types and Leiden subclusters through multiple analytic platforms. In Fig3A, we show select pathways that were found on multiple analytic platforms (Fig3A). Repeatedly results demonstrated consistency with parent cell type but began to elucidate the diversity of transcriptional responses. For example, moDerived macrophages had upregulation of antigen processing and presentation that was enriched in Cluster 1 along with IFN-γ signaling and oxidative phosphorylation enriched in Cluster 2. Cluster 7 also demonstrated an upregulation of antigen presentation as well as an anti-inflammatory response with *Sod2*, *Saa3*, and *C1qc*. Transitioning cells showed upregulation of phagocytosis enriched in Cluster 0 translation in Cluster 3. The ‘Notch-High’ cluster separated from the two main clusters of monocyte-derived cells, markedly denoted by a higher ratio of H37Rv-infected cells (63% as compared to 50% in moDerived macrophages, p<0.05; Fig2E,F). Likely due to the elevated inflammatory response, few transcriptional markers delineating the origin of cell type remained and Notch signaling was repeatedly enriched in multiple analyses, with key genes such as *Notch2, Cdkn1b*, *Jag1, Adam10, Dtx3l*, highly expressed. Alveolar macrophages clustered apart as well, but with common pathways of lipid metabolism although cluster 9 had upregulation of IFNγ response.

**Figure 3.**
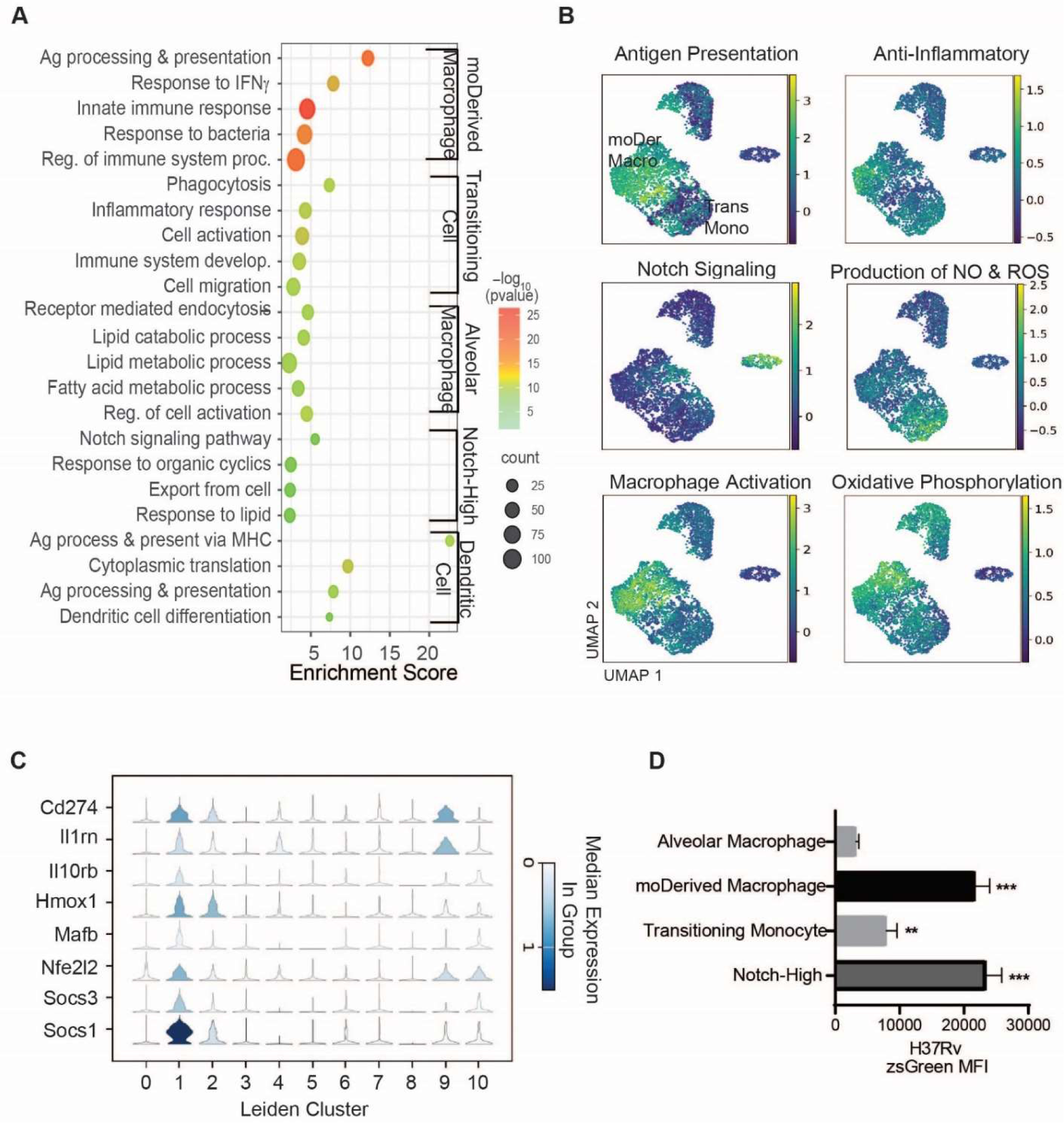
Recruited MNPs during Mycobacterium tuberculosis lung infection include a subset of cells with an anti-inflammatory signature. **A)** Differentially expressed genes between cell types were used for gene ontology analysis by ShinyGOv0.77, where color of dot denotes the negative log of false discovery rate (FDR) and size of the dot is the number of genes enriched in the denoted signature. **B)** Gene sets were constructed to score the likelihood of a particular cell with transcriptional enrichment of the denoted pathway using the average expression of associated genes (methods). Lighter color denotes higher expression of gene set and thus predicted attribute. moDer Macro = moDerived macrophage (leiden clusters 1, 2, 7) and Trans Mono = transitioning monocyte (Leiden clusters 0, 3,10). **D)** Violin plot of differentially expressed genes from the ‘anti-inflammatory’ pathway (SuppTable5) as delineated by Leiden cluster. **E)** ZsGreen MFI of infected MNP subsets from H37Rv/ZsGreen-infected mouse lungs (28 dpi).

To better visualize cellular heterogeneity, we utilized gene enrichment scores to delineate phenotypic predictions. Scores were developed utilizing genes our experimental model in conjunction with previously defined and associated signatures^54^ (Fig3B, SuppTable2, Methods). Fig3B shows the enrichment of multiple genes involved in the production of nitric oxide and reactive oxygen species in transitioning monocytes, antigen presentation and macrophage activation in moDerived macrophages, and notch signaling upregulation in the termed population. Remarkably, when compared to AM there is an anti-inflammatory signal concentrated in moDerived macrophages Cluster 1 from mice infected with H37Rv. Further, upregulation of key anti-inflammatory genes including *Hmox1,*^55^ *Il10,*^56^ *Il1rn* (Il1 receptor antagonist),^57,58^ and *Nfe2l2* (Nrf2)^2^ suggest that recruited moDerived macrophages could be more permissive for MTB replication (Fig3C). Indeed, infected moDerived macrophages harbor more bacterial burden (indicated by MTB fluorescence) than infected AM (Fig3D), consistent with previous findings that moDerived cells contain more live bacteria.^18^ Collectively, these findings suggest that virulent MTB recruits permissive macrophages with anti-inflammatory signatures despite the development of adaptive immune responses.

### ESX-1 is vital for the transcriptional induction of both recruitment and maturation of macrophages of infected lungs

Given that the lack of ESX-1 activity resulted in significantly less moDerived macrophages in MTB-infected lung, we analyzed recruited transitioning monocytes to understand if they behaved differently in the two infected environments. Compared to transitioning monocytes from uninfected mice, very few genes were differentially expressed in the same cell type from H37RvΔEccD_1_-infected mice. However, significant upregulation was noted for several genes involved in MHC presentation from mice infected with H37Rv compared to H37RvΔEccD_1_ (Fig4A,B). Additionally, Lars2 was upregulated in transitioning monocytes from H37Rv-infected mice compared to H37RvΔEccD_1_-infected mice. This gene is important for protein synthesis,^59^ but a direct correlation to antigen presentation remains unclear.^60^

**Figure 4.**
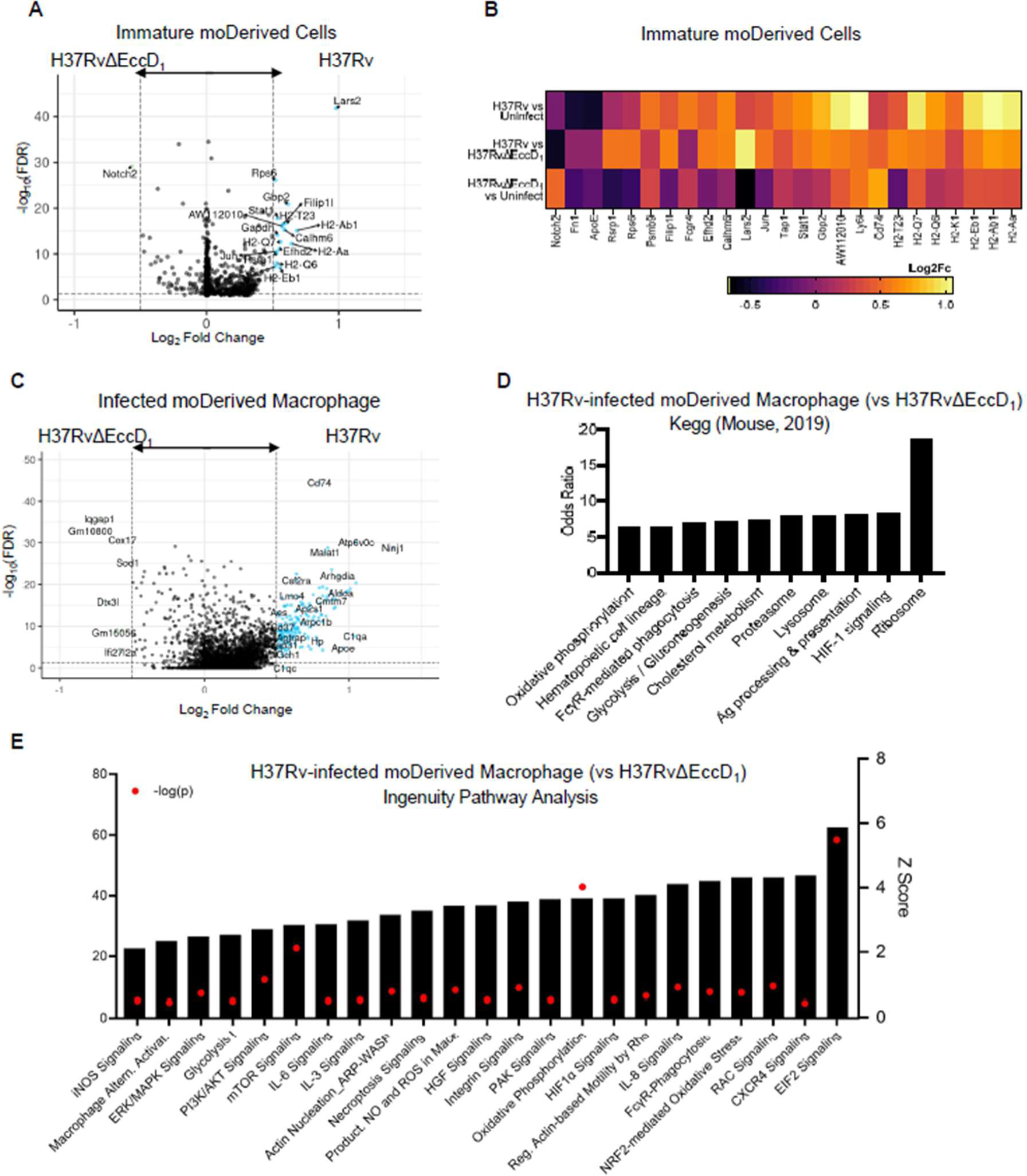
ESX-1 promotes macrophage activation and immunosuppressive response. **A)** Volcano plot of immature moDerived cells from mice infected with H37Rv and H37RvΔEccD_1_ analyzed for differential gene expression differences using MAST. **B)** Log2 fold change differences of differentially expressed genes between H37Rv versus H37RvΔEccD_1_ or each condition to cells present in the uninfected control mouse. **C)** Volcano plot of differentially expressed genes in mature monocyte-derived (moDerived) macrophages infected with H37Rv as compared to H37RvΔEccD_1_. **D)** Odds ratio for pathway enrichment using KEGG (mouse from 2019), with top 10 statistically significant pathways shown, removing redundancy. **E)** Qiagen Ingenuity pathway analysis was also queried using differentially expressed genes determined from MAST analysis with an FDR <0.1. Pathways shown are upregulated, with Z score on right X axis. The −log(FDR) is shown by red dots with value depicted on the left X axis.

To further understand ESX-1’s role within macrophages in the mouse lung, we directly compared H37Rv and H37RvΔEccD_1_-infected (i.e. zsGreen positive) moDerived macrophages using MAST to reduce bias by adjusting gene expression rate.^61^ Of note, this cellular population was not detected in the uninfected mouse, and therefore, directly comparing to cells not exposed to any bacilli was not feasible. This revealed 8 genes with a significant log_2_ fold change of ≥0.50 in H37RvΔEccD_1_-infected moDerived macrophages: 2 predicted or long coding genes (*Gm10800* and *Gm15056),*^62,63^ IFN inducible *Ifi27l2a*, as well as *Iqgap1*, *Dtx3l*, *Mtmr2*, *Cox17*, and *Sod1* (Fig4C). Among these, *Iqgap1* positively regulates autophagy and negatively modulates the type 1 IFN signaling pathway,^64,65^ suggesting a mechanism for better control of Mtb. Conversely, in cells infected with H37Rv, upregulation of 170 genes were noted (FDR <0.05) including key genes involved in HIF-1 signaling (*Hmox1, Nos2, Eno1*), classical antigen presentation (*Cd74, H2-Q7, H2-Dmb1*), and stimulation of cell death pathways (*Coro1a, Bax, NfKb2, Casp8, Hmox1*) (Fig4D). Qiagen Ingenuity Pathway analysis also revealed pathways favoring MTB survival,^2^ including alternative macrophage activation, TGF-β Signaling, CXCR4 signaling, NRF2 oxidative response, necroptosis, and pyroptosis signaling (Fig4E) and gene ontology concurrently supported upregulation of cellular locomotion and activation in H37Rv-infected cells. Together, the induction of an anti-inflammatory phenotype in Mtb-permissive macrophages, at least partially activated through multiple immune suppressive pathways (i.e., NRF2 oxidative response),^66–68^ is ESX-1 dependent.

### ESX-1 suppresses the innate inflammatory response early in intracellular infection

Bone-marrow derived macrophages (BMDM), derived from bone marrow progenitors and differentiated in vitro, have been utilized as primary cells to study MTB infection. To compare with transcriptional differences identified in vivo, BMDM derived from healthy C57BL/6 mice were infected for 24 hours with MTB expressing dsRed with or without functional ESX-1 through lack of the entire RD1 operon^69^ (Fig2A). Live cells were sorted on dsRed expression, multiplexed using lipid-tagged indices (MULTI-seq), and single-cell transcriptomic libraries were produced using 10X chemistry^70^ (Fig5A, SuppFig5A).

**Figure 5:**
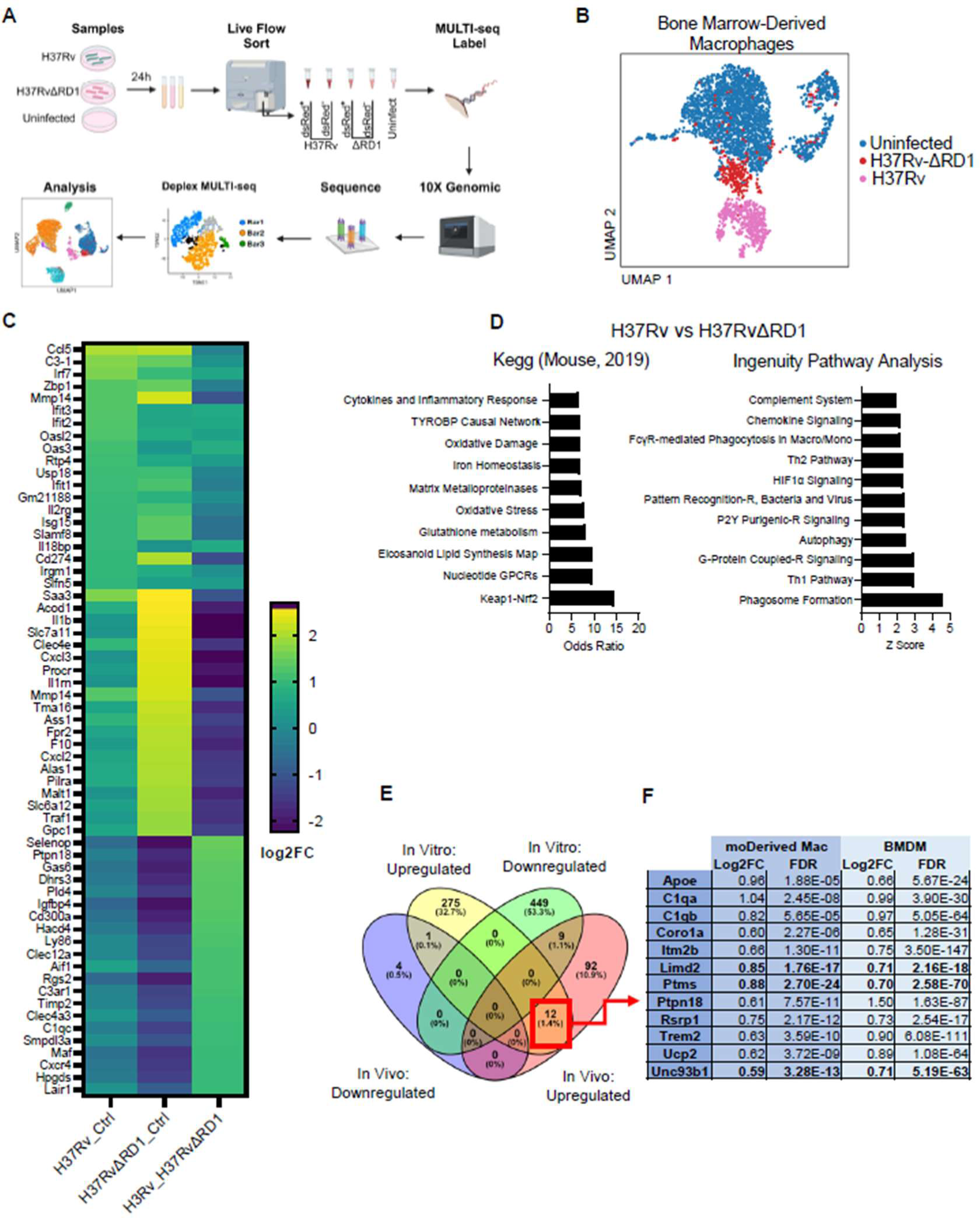
ESX-1 aids in the suppression of the inflammatory response in bone marrow derived macrophages. **A)** BMDM derived from C57BL/6 mice were infected with H37Rv or H37RvΔRD1 expressing dsRed for 24 hours, stained for viability, and flow sorted on dsRed expression. Cells were multiplexed and transcriptional libraries obtained through 10X chemistry. **B)** Dimension reduction of transcriptional libraries of each condition. **C)** Heatmap of the top 20 differential gene expressions by log2 fold change of each comparison: H37Rv or H37RvΔRD1 versus control and H37Rv versus H37RvΔRD1. **D)** Pathway enrichment of differentially expressed genes of BMDM infected with H37Rv as compared to H37RvΔRD1. **E)** Comparing differentially expressed genes up or downregulated in ESX-1 active or inactive strains from in vivo (moDerived macrophages) or in vitro (BMDM) **F)** revealed 12 genes concurrent in differential expression.

Dimension reduction demonstrated a clear separation of H37Rv-infected BMDM from uninfected cells, with H37RvΔRD1 separated but more closely related to the uninfected population (Fig5B). Overall, when compared to cells not exposed to bacteria, both H37Rv and H37RvΔRD1 infected cells resulted in an overall transcriptional upregulation of cytokine signaling, oxidative stress, and necroptosis (Fig5C). In the setting of an infection period of 24 hours, there was a surprising overlap in transcriptional changes between both infection states as compared to control. Nonetheless, differential regulation of key pathways was noted, such as ferroptosis and activation of the Th-1 pathway, which was significantly upregulated with the presence of ESX-1 (SuppFig5B).

To delve further into transcriptional differences between infection states, we directly compared BMDM infected with H37Rv with and without functional ESX-1. This short infection period did demonstrate the anticipated macrophage control of the intracellular bacteria through upregulation of complement genes, *Cxcr4*, *Aif1*, *Maf*, and pathway enrichment of oxidative stress, phagosome formation, and HIF1α signaling. However, differential gene expression also revealed upregulation of key genes associated with an anti-inflammatory response, such as *Selenop,*^71^ *Lair1,*^72^ and *Gas6,*^73^ with concurrent upregulation of the NRF2 pathway (Fig5C,D).

Different environmental factors, cell types, methods of analysis, and time course of infection are but a few factors that have long accounted for the inability to translate findings from cells infected with MTB in vitro to those in vivo. Using scRNAseq, we next analyzed what could be compared in our two models when ESX-1 function is lacking. In both lung moDerived macrophage and BMDM, 12 genes were consistently differentially upregulated in cells infected with H37Rv as compared to strains lacking functional ESX-1 (Fig5E,F). Multiple genes upregulated in both conditions have previously been demonstrated to play key roles in tuberculosis disease (*Apoe*^74,75^), immune evasion (*Trem2*^76,77^), intracellular survival (*Coro1a*^78^), biomarker discovery (*C1q*^79–81^), and general IFN-induced or inflammasome induction (*Itim2b*, *Ucp2)*. Three novel genes - *Limd2, Ptms*, and *Unc93b1 -* were also identified to correlate in MTB-cellular infection: Moreover, a specific reduction of recruited macrophages was noted in both models. Together, ESX-1 induces anti-inflammatory responses in macrophages in vitro and in vivo, which is associated with successful MTB intracellular survival/replication.

### Activated and mature monocyte-derived macrophages localize to the site of H37Rv bacilli

We identified heterogeneous cell types recruited to the lung 28 days after MTB infection. However, the spatial relationship of cells of each subset and bacilli cannot be determined with analysis of dissociated tissue. We therefore turned to spatial analysis of our identified cell types to elucidate if cellular maturation occurs at sites near MTB expressing ESX-1. Here, we utilized Vizgen MERSCOPE Platform (MERFISH),^82^ a spatial transcriptomics technique that combines highly multiplexed single-molecule fluorescence in-situ hybridization (smFISH) and labeling of specific proteins to provide subcellular transcript location, cell type clustering, and spatial analysis of unique cells of interest.^83^ For this, we optimized specific lung processing to retain high-quality RNA and co-labelled mycobacterium bacilli to provide a single image acquisition of infected versus bystander cells (Methods, SuppFig6).

Lungs from C57BL/6 mice infected with H37Rv for 28 days as in the previous experiments, and were labeled with a 122-gene panel constructed using gene signatures identified from our single-cell transcriptional atlas (Fig6A). We identified distinct cellular aggregates, previously described as non-necrotic lesions, further confirmed by MTB immunofluorescent staining.^84^ Using a high threshold to exclude background, infected cells were demarcated, and transcripts were analyzed for proximity to infected cells within, as well as clusters outside, the lesions (Fig6B, SuppFig6). Using data from eight distinct tissue sections from four different experiments, we identified 42 genes that were consistently upregulated in expression within 100 microns of MTB-infected cells (p<0.01 compared to blank probes; Fig6C). Transcripts concentrated within cells near those infected with H37Rv included *Hmox1*, *Nos2*, and *Hif1α*. As Hmox1 has been shown to inhibit Nos2 function,^85,86^ this suggests an anti-inflammatory signature in the lesion of C57BL6 mice. Multiple IFN-induced genes were noted to increase toward MTB, such as *Irf1*, *Nfkb2*, *Il1b*, and *Ifi30.* Still, the differentiation of type-I and II could not be definitively made with the available probe signatures. In addition, the transcriptional upregulation of MHC-II antigen presentation did increase toward MTB-infected cells (*Cd74*, *H2Ab1*, *H2Eb1*), as well as an increase of *C1qb* and *C1qbc* concordant with the signature of the mature moDerived macrophage seen in our transcriptional atlas. Indeed, the moDerived macrophage gene signature was increased, and conversely transitioning monocyte signature decreased, in proximity to cells actively infected with MTB (Fig6D).

To ensure these genes and others of interest were expressed by MNPs, we first confirmed that *Cd74* was expressed by over 90% of cells. Secondarily, we utilized a gene signature that included nine canonical markers for monocytes and macrophages with no detectable expression in other mouse lung cells in a published single-cell dataset^87^ (*C1qb*, *C1qc*, *Clec4a2*, *Dnase1l3*, Ms4a7, *Sell*, *Trem2*, *Mmp9*, *Clec4e*) and found over 80% of the cells analyzed expressed this signature, with the highest expression found in infected and neighboring cells (Fig 6E, SuppFig6C). Re-analyzing gene expression patterns in the population of strictly defined MNPs expressing detectable levels of these markers validated the up-and down-regulation of genes shown above, including Nos2 and Lyve1 (Fig6F and SuppFig6D-E).

**Figure 6:**
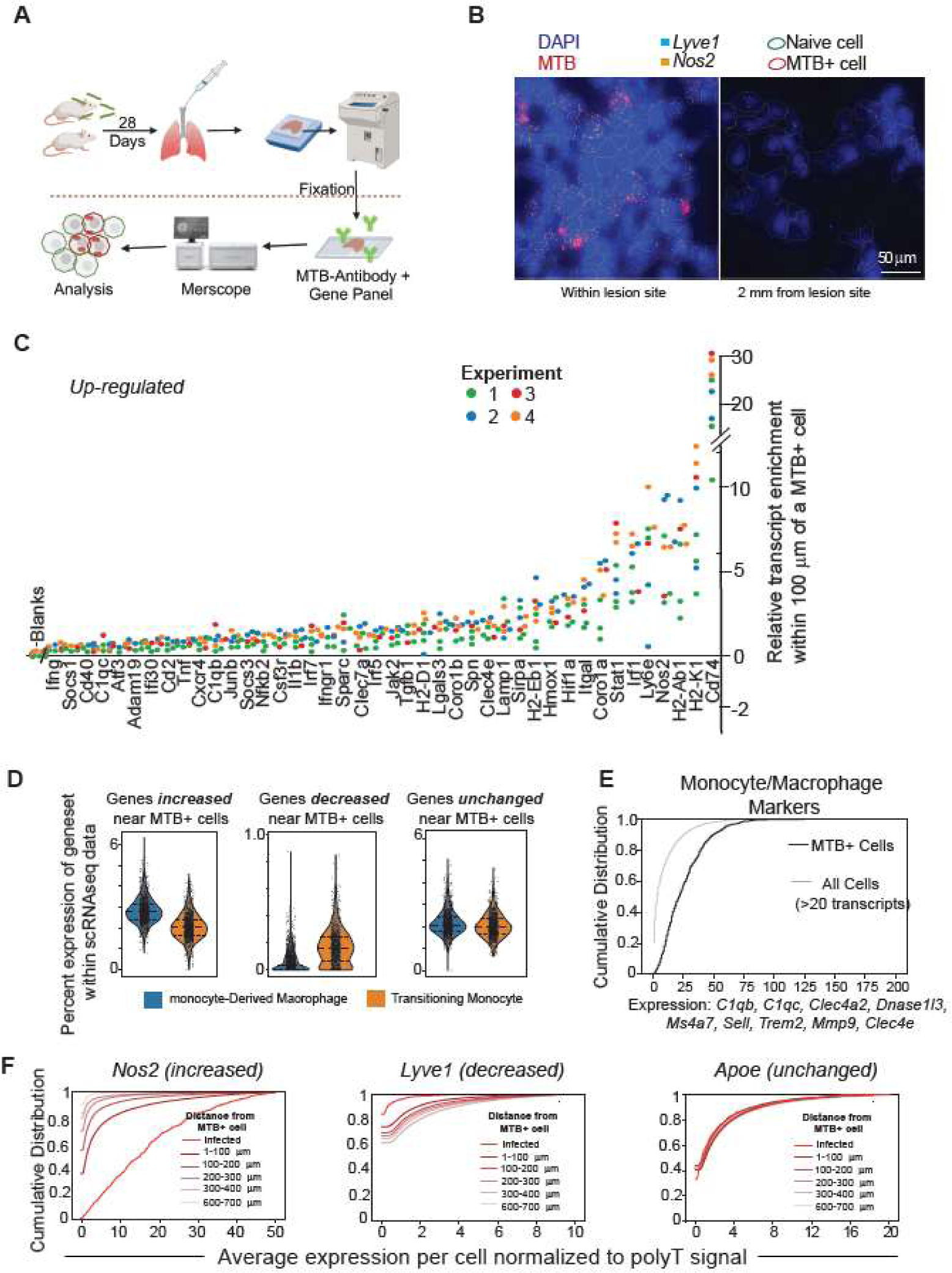
MTB induces maturation of macrophages only in close proximity to its presence. **A)** C57BL/6 mice were infected with H37Rv for 28 days, and after perfusion with PBS, lungs were inflated and freshly frozen in OCT. 10 µm slices were fixed and processed for Vizgen Merscope visualization of both MTB (using Ab905) and gene panel (SuppTable9). **B)** Representative images from Merscope Visualizer demonstrating MTB (red), DAPI (blue), and cell boundary segmentation with red as meeting threshold for MTB infection (i.e. MTB+) and green under the threshold (i.e. naïve). All circled cells express at least 20 transcripts from the panel. *Lyve* 1 and *Nos2* are noted by blue and orange dots, respectively. **C)** Plot showing transcripts that are significantly upregulated within 100µm of MTB-infected cells, with each dot representing an experiment. Y-axis is the difference between mean expression for a transcript in cells within 100µm of MTB-infected cells as compared to the mean expression of transcripts within all cells normalized to polyT (total mRNA per cell). All genes had a p-value of <0.01 compared to pooled blank transcripts. **D)** Violin plots showing the percent expression of upregulated (shown in C), downregulated (*Lyve1, Sup1, Scarf2, Itga6, Jun, Icam2, Car4*), and unaffected genes within SmartSeq atlas shown in Figure 4, with quartiles distinguished by lines within the violin plot. **E)** Cumulative expression of representative markers for monocyte or macrophages. X-axis represents the normalized gene counts from all listed MNP genes combined, and the Y-axis is the percentage of cells expressing less than or equal to the given expression level. <1% of infected cells (black line) and <20% of all cells (grey line) have no detectable expression of the marker genes. **F)** Representative cumulative distribution plots showing a gene highly enriched near MTB-infected cells (*Nos2*), downregulated near MTB-infected cells (*Lyve1*), and one that is unaffected by MTB presence (*Apoe*). Each color represents a bin of width 100 µm. Of note, not all lines start with y=0 due to sparse gene expression levels.

The gene signature downregulated in neighbors of MTB-infected cells was less robust than the unregulated signature, with only seven genes consistently downregulated. One of these genes, *Lyve1*, is expressed in both endothelial cells and perivascular anti-inflammatory macrophages^37^. While the analysis shown here may include some endothelial cells, Lyve1 was downregulated even in strictly defined MNPs (SuppFig6D-F), in contrast to other anti-inflammatory genes including *Mafb* and *Hmox1* which were consistently increased in expression near sites of infection. Therefore, while cells with an anti-inflammatory signature are recruited towards infected macrophages, they do not appear to be the Lyve1-high cells described elsewhere.^88^

We asked what signature difference occurs between an infected macrophage and close proximity bystander monocyte or macrophage. Here, only strictly defined MNPs with at least five counts of the MNP markers were analyzed to compare similar populations. As shown in figure 7, *Nos2* was the most significantly differentially expressed gene in the infected versus bystander cells (Fig7A). Conversely, genes related to MHC-II expression were downregulated in infected cells as previously described.^89^ Although limited by the gene panel, differential expression in the spatial analysis correlated with the comparison of infected to bystander moDerived macrophages and transitioning monocytes in the dissociated tissue analysis with significant upregulation of *Nos2* and downregulation of *Coro1a* (Fig7B). Pathway prediction model could not be completed but the enrichment of key anti-inflammatory genes warrants further evaluation. Overall, this novel technique supports that mature moDerived macrophages were concentrated near the bacillus-infected cells, with active differentiation in close proximity.

**Figure 7:**
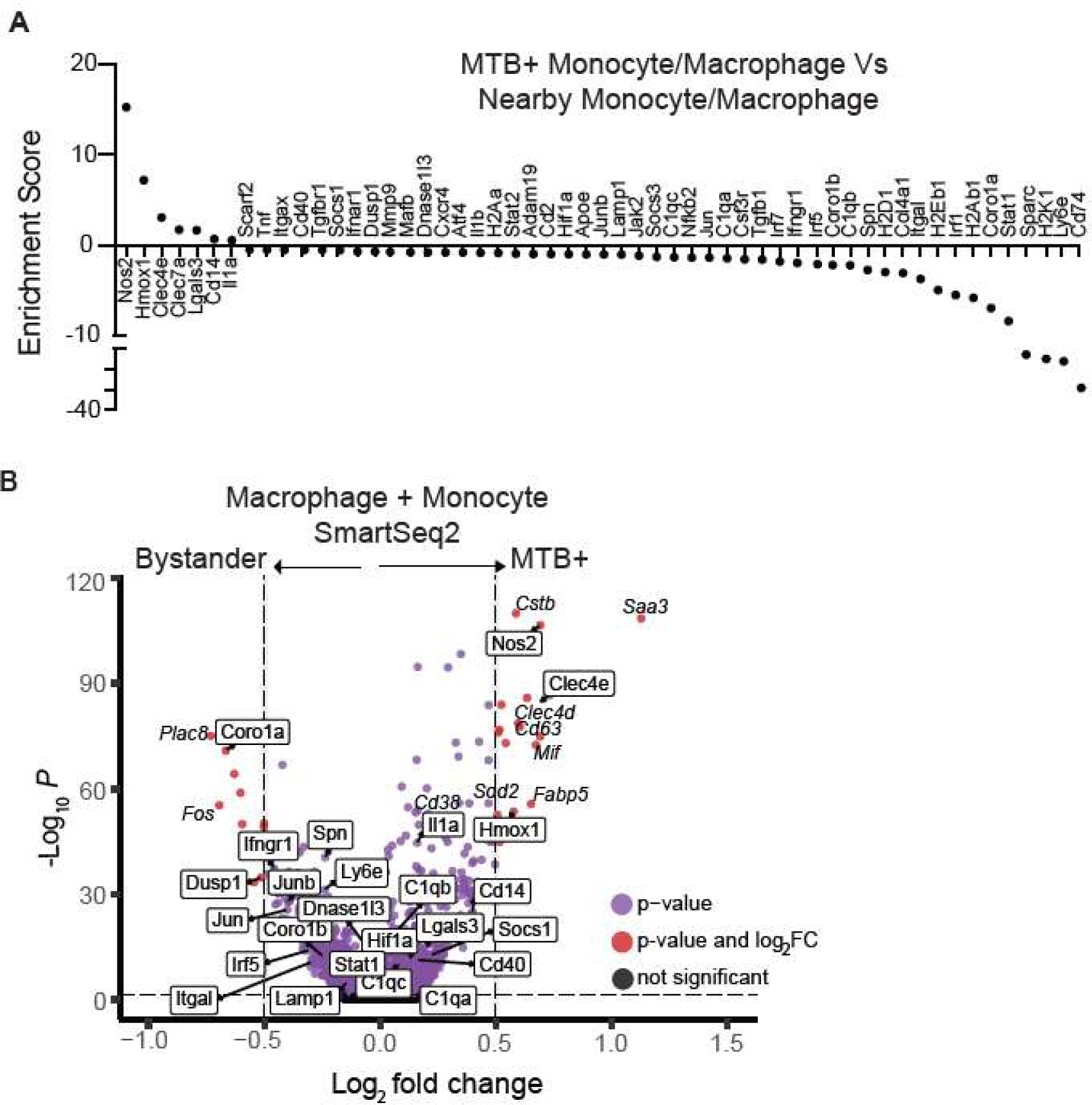
Few key differentially expressed genes between MTB-infected and bystander cells persists with tissue dissociation while the majority are diminished. **A)** Gene enrichment score (difference in average normalized gene expression) between infected monocytes and macrophages as compared to neighboring (within 100 µm) uninfected cells. All cells included in the analysis expressed 5 or more transcripts from a list of canonical markers not expressed in other cell types (*C1qb, C1qc, Clec4a2, Dnase1l3, Ms4a7, Sell, Trem2, Mmp9, Clec4e*). All genes displayed had a p-value of <0.001 based on a permutation test with 1000 iterations. **B)** Differential gene expression of moDerived macrophages and transitioning monocytes from SmartSeq2 by MAST, comparing cells expressing MTB fluorophore (MTB+) versus those not expressing fluorophore (bystander) from H37Rv-infected mice. Genes from panel A are boxed for visualization to compare between the two methods.

## Discussion

In this study, we demonstrate that the heterogeneity of recruited MNPs is significantly altered in the absence of ESX-1 function, and the priming of immature recruited MNPs to an anti-inflammatory transcriptional phenotype differs when ESX-1 is active. Further, spatial transcriptomics provides understanding of gradient macrophage activation at the angle of individual cell infection. Our study, which focuses on a specific time point after the adaptive immune response has been well developed, used state of the art techniques to investigate the role of ESX-1 in heterogeneous cellular recruitment and elucidates the transcriptome within spatial contexts as they relate to MTB bacilli. These results demonstrate how novel tools can be integrated to enhance our understanding of MTB’s ability to drive the maturation of macrophages for its benefit.

The diversity of steady-state myeloid cells within the lung is now gaining recognition but remains incompletely understood,^32,34,90,91^ complicating our ability to describe the innate immune response during later stages of infection. This is exemplified by our observation that the ImmGen reference datasets cannot provide refined cell identification in the setting of tuberculosis infection. Indeed, we previously described two major MNP subsets through single-cell flow cytometry analysis,^14,17,18,26^ and as anticipated based on recent evidence of macrophage heterogeneity,^32–34,54^ these cellular populations exhibit transcriptional heterogeneity. A recent publication has also revealed heterogenous monocyte and macrophage responses during MTB infection at an earlier time during infection.^92^ Our work extends these findings to a period when the adaptive immune response is fully activated. Additionally, using SmartSeq2 provided an expanded transcriptional library and the ability to capture rare events of cells infected with an attenuated MTB strain.

Historically, understanding of the macrophage response to infection and inflammation has relied on a dichotomous view of M1 and M2-like macrophages.^93–95^ However, recent robust data demonstrates that macrophages exhibit a much more diverse response, further substantiated by the explosion of single-cell transcriptional analysis.^30,54,96^ Our work aligns with this heterogeneity, confirming variation within, and transition states between, what might be considered M1 or M2-like states. Further, modeling of such states also does not occur in vitro.^97^ We therefore refrained from dichotomizing our clusters to avoid oversimplification that could lead to misinterpretation. In this paradigm, a recurrent theme of anti-inflammatory induction is observed during H37Rv infection. Indeed, others have found significant anti-inflammatory signals in recruited cell subsets at an earlier time point before adaptive responses to MTB have been well developed.^92^ Our work extends this with additionally demonstrates that the recruitment of such cells is ESX-1 dependent. Further, our findings reveal the plasticity of macrophage responses during MTB infection.

ESX-1 is important in the inhibition of phagolysosome fusion.^38,40^ Additionally, it is involved in phagosome perforation,^98^ facilitating the direct escape of MTB to the cytosol. Membrane disruption through ESX-1 activity^99^ can ultimately lead to activation of Il-1β and necrosis^100,101^ or pyroptosis.^102^ Release of the bacillus can happen within a short time after infection^99,103,104^ making initiation in a primed and habitable cell type beneficial for the bacterium’s survival. In our study, we observed a significant upregulation of *Lars2* in recruited monocyte-derived cells only in the presence of active ESX-1. Lars2 encodes mitochondrial leucyl-tRNA synthetase and likely protects against mitochondrial stress through boosting mitochondrial translation,^105^ and is upregulated during MTB infection when there is experimental induction of mitochondrial stress.^106^ Therefore, future studies are needed to test if MTB can induce this pathway in bystander-recruited cells as a survival tactic to avoid apoptosis during infection and allow intracellular persistence.

Macrophages were recruited at a higher proportion compared to monocytes in H37Rv infection compared to H37Rv lacking functional ESX-1. While this could be partially explained by different dynamics of infection,^7,37^ the differential responses of macrophages both in vivo and in vitro provides further insight. Within both models, functional ESX-1 induces multiple immunosuppressive pathways. For example, NRF-2 signaling, considered to have anti-inflammatory defense through the expression of various antioxidants,^107,108^ has been shown to be an important component of alveolar macrophage’s inability to control intracellular MTB.^2^ NRF2 also induces HMOX1 to inhibit NOS2-mediated nitric oxide production, which depends on ESAT-6 secreted by ESX-1^68^. In addition, alternative activation pathways (e.g. IL-10)^109–111^ and TGF-beta signaling^112,113^ have been shown to not only inhibit the anti-MTB activities of macrophages but also suppress lung T cell activation, suggesting that anti-inflammatory responses suppress both innate and adaptive responses to support MTB persistence in recruited macrophages during chronic infection. In our data, recruited macrophages demonstrated induction of oxidative stress and glycolysis, but simultaneously exhibited an ESX-1 dependent upregulation of immunosuppressive pathways both in vitro and in vivo. This observation holds in a spatial context, where cellular maturation and IFN stimulation occurred alongside the upregulation of key protective factors. We hypothesize that ESX-1 assists in the recruitment of cells primed for manipulation and MTB persistence, and further work is aimed at understanding the signals required for recruiting this subset from the bone marrow.

Much of what is known about cellular permissibility to MTB and ESX-1’s role in phagosome release has been conducted in vitro, often using self-propagating cell lines such as THP-1. In our study, while some general pathways did show correlation in vivo and in vitro, only a handful of specific genes aligned in both models. This discrepancy is not novel,^114^ especially to the field of MTB infection. However, our work underscores the importance of employing innovative methods to bridge in vivo and in vitro models and allow for downstream mechanistic investigations to reduce animal use. Notably, our comparative transcriptomic analysis has identified 3 potential genes of interest. *Limd2* is previously shown to be involved in cell motility,^115–117^ *Ptms* implicated in chromatin remodeling,^118^ and *Unc93b1* instigated in immune regulation and macrophage recruitment during viral infection^119^ are individually targets of interest. Importantly, these findings demonstrate the importance of comparative genomics to combine publicly available data to elucidate novel factors important in MTB pathogenesis and potential diagnostic markers of infection.

A key to comprehending the intricate interplay of MTB with recruited, infected, and bystander cells lies in elucidating spatial interactions. Analyzing this interaction at the transcriptional level offers an in-depth understanding of the microenvironment. We have previously shown that Vizgen MERSCOPE, a microscopy-based spatial transcriptomic method,^82^ provides equivalent cell type resolution and enhanced sensitivity compared to standard scRNA sequencing methods.^83^ Using a panel of 122 genes, we identified cell types described in our 10X and SmartSeq2 atlas in space, as well as unveiled genes that exhibit differential regulation across the myeloid microenvironment more comprehensively than previously reported.^120,121^ MHC class II genes and Nos2 were the most significantly upregulated transcripts toward the MTB-infected cell.^120^ Surprisingly, we observed a reversal in the expression of *Lyve1*, even though macrophages expressing high levels of Lyve1 have been implicated in an anti-inflammatory response and are associated with blood vessels in the lung.^88^ This discrepancy is likely due to the turnover of recruited cells and highlights how chronic infection induces distinct macrophage responses that do not correlate with other inflammatory responses. However, other markers of an anti-inflammatory response did increase toward the MTB-infected cell and are congruent with our findings that MTB infection induces the recruitment of anti-inflammatory macrophages. Spatial analysis is also instrumental in fully understanding transcriptional differences of infected and bystander cells in the microenvironment, as noted by the fact that differential gene expression is diluted when analyzing the entirely dissociated tissue. Ontogeny analysis is beyond the scope of this work but will be insightful to elucidate where these more permissive cells originate and points of transition in conjunction with this approach.

There are a few notable limitations to these experiments. Here, we used SmartSeq2 which, while highly effective, resulted in a restricted number of cells due to its inherent limitation in yield. However, the high-quality libraries and capability to capture rarer events of H37RvΔEccD_1_-infected mitigated this limitation. In fact, we found lower quality libraries in our 10X library than SmartSeq2 due to longer processing time from sort to load, increased fragility of cells after cell sort and washing, and ability to capture rarer events immediately at sort.^122,123^ This also provided the ability to capture discrete subsets such as Notch-high subset, which contained high MTB fluorescent expression, suggesting larger bacilli per cell. Further, this work is largely transcriptional and differential gene expression does not always correlate with significant differences in protein expression warranting further investigation into expression variation among cellular subtypes. Despite these limitations, the recurrent and consistent findings obtained in this study are noteworthy. Our study is limited to a specific time point post-development of the adaptive immune response, providing only a snapshot of the macrophage response to MTB without ESX-1, as the dynamics of infection are delayed during attenuated bacterial infection. Leveraging computational approaches, we believe our work can contribute to the growing body of research utilizing single-cell transcriptomics to elucidate the dynamics of recruited macrophage responses over time and enhance the understanding of MTB survival in host MNPs.

In summary, our study underscores ESX-1’s vital role in the recruitment of specific anti-inflammatory MNP and promoting macrophage maturation. Our data suggest that MTB induces the recruitment of an immature cell type primed for its own survival. Spatial transcriptional analysis provides a valuable avenue for understanding the in-situ heterogeneity of gene expression in the infected microenvironment and offers further evidence of an induction of key protective factors for bacterial survival.

## Methods

### Mice and Care

C57BL/6 mice were purchased from the Jackson Laboratory and were between the ages of 6 to 8 weeks at the beginning of the experiment. For infections with *M. tuberculosis*, mice were housed under barrier conditions in an ABSL-3. Mice of both sexes were used. Mice were euthanized by CO_2_ asphyxiation followed by cervical dislocation. All experiments were performed with the prior approval of the University of California-San Francisco Institutional Animal Care and Use Committee (IACUC).

### Bacterial strains and growth

All MTB strains were derived from an H37Rv background and grown as previously described.^124^ Constitutive fluorescent protein expression was obtained by transforming with pMSP12::mCherry, pMV261::ZsGreen, or pMSP12::dsRed2, and maintained by culturing in the presence of 50 μg/ml kanamycin. H37RvΔEccD_1_ mutant was constructed via abortive transduction as a method of allelic resistance substrate (AES) delivery.^49,125^ AES were generated by flanking a hygromycin resistance cassette with ∼500bp of *EccD_1_* (*Rv3877*). Transduced bacteria were selected on hygromycin-containing agar, and confirmed through PCR, followed by Sanger sequencing of the region of interest.

### Aerosol infection

Bacterial strains were grown and stored for stock inoculum, with mice infected via the aerosol route using an inhalation exposure system (Glas-Col), as previously described.^7,14,17,18,26^ The target dose was 50-75 CFU/mouse for H37Rv and 500 CFU/mouse for H37RvDEccD_1_. The infectious dose was quantitated on day 1 by plating whole lung homogenates from 3 mice on Middlebrook 7H11 agar. The bacterial load at the time of analysis was also determined by serial dilutions plated on Middlebrook 7H11 agar. Colony Forming Units (CFUs) were counted after incubation of plates at 37**°**C for 3 weeks.

### Cell culture infection

BMDM were derived from bone marrow cells cultured in DMEM (Gibco 11965092), 10% heat-inactivated FBS (HI-FBS) and 20 ng/mL recombinant murine M-CSF (PeproTech) as described elsewhere.^7^ Before infection, cells were washed twice with PBS. Bacterial strains were grown without tween and stored as previously described.^125^ Stock was thawed and any remaining clumps removed by filtering through a 0.05uM filter. Cells were bathed in infection media at a MOI of 1 for 4 hours, washed, and media replaced. 24 hours post infection media placement, live cells were lifted gently, washed, and stained by Zombie Aqua Fixable Viability Dye (BioLegend, 423101) prior to flow sorting.

### Lung homogenate preparation, Flow cytometry and cell sorting

Lung homogenates were prepared as previously described with modifications.^18^ For SmartSeq2 experiment, mice were retro-orbitally injected with anti-CD45-PE for 2 minutes prior to euthanasia. For all experiments, after euthanasia, lungs were perfused with 10 mL of PBS/2 mM EDTA via right ventricle immediately after euthanasia. Lungs were chopped into small pieces, placed in digestion media containing 4 mL of RPMI-1640/5% HI-FBS containing 1 mg/mL collagenase D (Sigma-Aldrich) and 30 μg/mL DNase I (Sigma), minced with a gentleMACS (Miltenyi, lung program1), and digested for 30 min at 37 °C. Tissue was further minced with the gentleMACS (lung program2) and then passed through a 70-μm cell strainer, rinsed, and red blood cells lysed with 3mL ACK lysis buffer (Gibco) for 3min and washed twice with RPMI-1640/5% HI-FBS.

Cells were stained with Zombie Aqua Fixable Viability Dye (BioLegend, 423101) for 15 minutes at 4°C, washed, and blocked at 1:100 with CD16/CD32 (BD, 553142) for 10 minutes prior to appropriate antibody staining for 30 minutes as below, diluted in Brilliant Stain Buffer (BD, 566349).

| **Antibody** | **Clone** | **Source** |
| --- | --- | --- |
| CD11c-BV605 | N418 | BioLegend 117334 |
| CD11b-BV711 | M1/70 | BioLegend 101241 |
| Ly6G-BV421 | 1A8 | BioLegend 127628 |
| Ly6C-PE-Cy7 | HK1.4 | BioLegend 128018 |
| I-A/I-E(MHCII)-PE-Dazzle | M5/114.15.2 | BioLegend 107647 |
| SiglecF-APC-Cy7 | E50-2440 | BD Biosciences 565527 |
| CD90.2-PE-Cy5 | 30-H12 | BioLegend 105314 |
| CD19-PE-Cy5 | 6D5 | BioLegend 115510 |
| NK1.1-PE-Cy5 | PK136 | BioLegend 108716 |
| CD45-PE | 104 | BioLegend 109808 |
| CD45.2-PerCp/Cy5.5 | 104 | BioLegend 109827 |

Fluorescently labeled live cells were then acquired using a BSL3-contained Sony MAC900 cell sorter as shown in supplementary figures. For flow cytometry, acquisition data was analyzed using FlowJo software (TreeStar). To obtain mean fluorescent intensity at time or cell sort, data was reviewed using the Sony MA900 software from sort for each plate run, and using Boolean expression, tabulated for integration into the transcriptional data object.

### Single cell cDNA Generation, Library Construction and Sequencing

#### SmartSeq2

Single cells were sorted and RNA transcriptomics obtained following the Smart-seq2 was modified from Picelli et al,^46^ using high-throughput liquid dispensers detailed elsewhere which includes: preparation of lysis plates; cDNA synthesis, library preparation using Illumina Nextera XT kit, library pool and ampure bead clean-up; and quality control. To provide feasibility within a BSL3, plates containing RT were modified to provide centrifugation for distribution. For further details, please refer to https://dx.doi.org/10.17504/protocols.io.8epv5xx54g1b/v1. Library pools were sequenced on the NovaSeq 6000 Sequencing System (Illumina) using 2 x 100-bp paired-end reads and 2 x 12-bp index reads with a 300-cycle kit (Illumina, 20012860).

#### 10X Single Cell 3’ GEM

For lung myeloid cells, single cells were sorted as described, washed, manually counted with trypan blue on a hemaocytometer. Using a 3’ v3.1 kit according to the manufacturer’s protocol, samples were loaded on a 10X Genomics Chip A without multiplexing with each condition loaded on 2 separate lanes to obtain GEMs (Gel Beads-in-emulsions),). For in vitro bone marrow derived cells, after single cell sorting, washing and manual counting, single cells were first labeled using the MultiSeq labeling method as detailed further in https://doi.org/10.1038/s41592-019-0433-8 prior to using a 3’ v3.1 kit and loading on a Genomics Chip A for GEM construction. cDNA and cleanup were done following manufacturer’s protocol (10X).

#### Library construction & sequencing

From generated cDNA in the previous step, libraries were constructed using the Chromium Next GEM Single Cell 3’ Library Kit v3.1. Library yields were assessed using Agilent Tapestation and quantified via qPCR (KAPA Library Quantification Kits for Illumina platforms, KK4828) on a CFX96 Touch Real-Time PCR Detection System (BioRad). Plate pools were normalized and combined at an equimolar ratio of 2 nM to make a sequencing library pool. Library pools were sequenced on the NovaSeq 6000 Sequencing System (Illumina) using 28 + 91 -bp paired-end reads and 8-bp Index 1 reads with a 100-cycle kit (Illumina, 20028400).

### Data Extraction & Analysis

#### Alignment

**-** Sequences from the NovaSeq were de-multiplexed using bcl2fastq version 2.20. Sequenced reads from SmartSeq2 libraries were aligned with Gencode v.M19 annotations and Gencode v.M19.ERCC using STAR version 2.5.2b with parameters TK. Sequenced reads from 10x Single Cell 3’ libraries were aligned with a custom mouse reference genome with added mCherry sequences that was created using CellRanger v.5.0.1 software.

#### Dimension reduction and differential gene expression

After standard preprocessing, dimension reduction was conducted using Scanpy (Python). Objects were converted to R for differential gene expression (DE) using MAST. Significantly expressed genes (FDR <0.05) were used for gene ontology (ShinyGOv0.61 and EnrichR). Ingenuity Pathway Analysis (Qiagen) was employed on genes with FDR <0.1 and ranked by log2fold change to identify regulation direction of pathways.

#### Gene enrichment scores

Enrichment scores were calculated from top enriched pathways and previously described macrophage transcriptional phenotypes using Scanpy^54,93^ (SuppTable2).

### Spatial Transcriptomics

C57BL/6 mice aerosol infected with H37Rv-zsGreen at ∼75 CFU/mouse for 28 days were sacrificed. Lungs were perfused with 10 mL of PBS/2 mM EDTA via right ventricle, followed by inflation through the trachea with 1mL of optimal cutting temperature (OCT) mixed with PBS at 1:1. Lungs were embedded in OCT, flash frozen, and stored at −80°C for later cryosectioning. To confirm quality, RNA integrity number (RIN)^126^ was obtained by extracting RNA from each block (QIAGEN RNeasy Mini Kit, Cat. No. 74104) and analyzing quantity on an Agilent Tapestation system. OCT-embedded tissue blocks were later cryosection at −20°C to a thickness of 10 μm and mounted onto MERSCOPE Slides (Vizgen, PN 20400001). The mounted tissues were immediately fixed with 4% PFA at 37°C for 30 minutes, washed with 3x PBS and stored in 70% ethanol at 4°C for no more than 1 mo before proceeding following the manufacture’s protocol (MERSCOPE) as previously described^83^ with the following exceptions. First, cell boundaries of mounted infected tissues including MTB bacterium were stained with the Vizgen Cell boundary Staining Kit (Cat. no. 10400009) in conjunction with a polyclonal MTB antibody (Abcam, ab905) raised in rabbit diluted to 1:100. After washing, the Secondary Staining Mix (Vizgen, PN 20300011) was diluted with a blocking solution in two step dilution (1st - 1:100, 2nd - 1:33) to ensure correct ratio of primary to secondary staining since a custom primary MTB antibody was used.

#### Imaging

The MERFISH gene panel consisted of 122 genes, designed by selecting top differentially expressed genes in each cell type from the myeloid scRNA sequence atlas, canonical markers, and genes of interest. The probes were hybridized, and samples washed prior to gel embedding without deviation from manufacter protocol. Samples were then treated with a clearing solution and imaged 1 day later per MERSCOPE User Guide 91600001. Only one gel-embedded and cleared sample was prepared per imaging run. The gel-embedded and cleared sample was washed prior to adding the DAPI / Poly T Staining Reagent, and again prior to assembling the onto the MERSCOPE Flow Chamber (Vizgen, PN 10300003). Once fully thawed, the MERSCOPE 140 Gene Imaging Cartridge was activated and a layer of mineral oil was carefully added on top of the imaging buffer. The assembled flow chamber with sample and imaging cartridge was placed in the MERSCOPE, image mosaic was acquired from the regions of interest, and imaging of the sample was performed with multiple rounds of hybridization of fluorescent probes which resulted in a raw images stack that contained about 1 TB of data.

#### Image Analysis

The full code can be found here: https://drive.google.com/file/d/1Z5UQohdkp0xEZXBprs127fkzqV7nuWCJ/view?usp=drive_link

All figures represent 2 mice analyzed in 4 separate experiments each measuring 1-3 lung sections. “Replicate” refers to an individual section. Results were highly concordant whether considering separate sections, experiments, mice, or the sum total.

For the spatial analysis showing differential abundance of transcripts near TB+ cells, the Vizgen cell by gene matrix was first filtered to include cells with at least 20 transcripts and with a volume between 200 and 5000 cubic micrometers and a cumulative DAPI signal greater than 2 million. Areas of tissue that showed evidence of damage (i.e. tissue folding or tearing) were excluded from downstream analysis. TB+ cells were defined as any cell whose top 100 brightest pixels in the anti-TB channel averaged above 300 units. Any cells with a standard deviation in this channel lower than 150 (that is, cells with uniform brightness) or further than 400 um from their two nearest neighbors (that is, isolated in space) were not counted as confidently TB+ cells. This threshold was chosen by picking a range of values, labeling the resulting positive cells in the Merscope Visualizer app, and checking for accurate labels compared to the TB antibody channel. The chosen thresholds limited the selection of clearly negative cells while positively labeling the majority of cells with three or more TB+ puncta, and labeled few cells in the negative control slices.

We then calculated the euclidean distance from each cell in the tissue to its nearest TB+ neighbor cell. Transcripts enriched near TB+ cells were identified by binning the cells found within 100 micrometers (∼5 cell widths) of a TB+ cell (excluding those infected) and comparing the average transcript expression within these cells to the average transcript expression in all cells 100 micrometers or further from an infected cell. First, each cell’s transcripts were normalized to the total RNA content of the cell by dividing the transcript counts by the cellular polyT signal and multiplying by the average polyT signal for that section. Expression levels for each transcript were then compared within each section using a bootstrap permutation test (run 10,000 times) comparing the actual difference in means to the difference in means when the cell location labels are randomly distributed. Transcripts with a mean increase or decrease in average count near TB+ cells that was greater than the expected (random) distribution in at least 9,990 out of 10,000 permutations (p<0.001) were counted as significant. The measurements for all replicates were combined and a Kruskal-Wallis test was run to find transcripts whose enrichment over all the replicates differed from the background (Blank) signal. The violin plots were generated by selecting genes that were consistently down-regulated (‘Lyve1’, ‘Dusp1’, ‘Scarf2’, ‘Itga6’, ‘Jun’, ‘Icam2’, ‘Car4’) across all replicates or up-regulated across all replicates (Kruskal-Wallis test vs. all blanks, p<0.01; [’Ifng’, ‘Socs1’, ‘Cd40’, ‘C1qc’, ‘Atf3’, ‘Ifi30’, ‘Adam19’, ‘Cd2’, ‘Tnf’, ‘Cxcr4’, ‘C1qb’, ‘Junb’, ‘Socs3’, ‘Nfkb2’, ‘Csf3r’, ‘Il1b’, ‘Irf7’, ‘Ifngr1’, ‘Sparc’, ‘Clec7a’, ‘Irf5’, ‘Jak2’, ‘Tgfb1’, ‘H2D1’, ‘Lgals3’, ‘Coro1b’, ‘Spn’, ‘Clec4e’, ‘Lamp1’, ‘Sirpa’, ‘H2Eb1’, ‘Hmox1’, ‘Hif1a’, ‘Itgal’, ‘Coro1a’, ‘Stat1’, ‘Irf1’, ‘Ly6e’, ‘Nos2’, ‘H2Ab1’, ‘H2K1’, ‘Cd74’]. These genes were grouped into an “upregulated”, “downregulated”, or “neither” geneset and their total percent expression was calculated for each cell in the scRNAseq dataset (Smartseq only; macrophage and dendritic cell populations) shown earlier in the paper.

## Statistical analysis

Flow cytometry experiments were performed at least three times. Results are expressed as mean and standard deviation, with unpaired t test with Welch correction and Holm-Sidak multiple comparisons, with a 95% confidence interval was used to compare experimental groups, with p<0.05 considered significant. Statistical approaches for gene expression are detailed in the above.

## Data Availability

Transcriptomic data will be available at GSE under accession number GSE263892. Code notebooks for Jupyter and R analysis are available on Github repository: https://github.com/bspeco/ESX1_macrophageheterogeneity_scRNA.git. The processed data for MERFISH are available upon request.

## Acknowledgments

We would like to thank Dr. Joel Ernst, the University of California San Francisco, for his extraordinary insight in the review of this manuscript.

## Author contributions

W.Z., A.P., O.R., N.N., B.S.Z. designed the research. O.R., N.N., B.S.Z. supervised the study. W.Z., M.B., A.S., R.A., S.S., N.A., B.S.Z. performed the experiments. B.S.Z., M.B. performed single cell transcriptional analyses and L.D., J.L., A.Z. performed spatial transcriptional analyses, with oversight by A.P. and N.N. W.Z., L.D. wrote the manuscripts with direct insight from all authors.

---

**Supplemental Figure 1:**
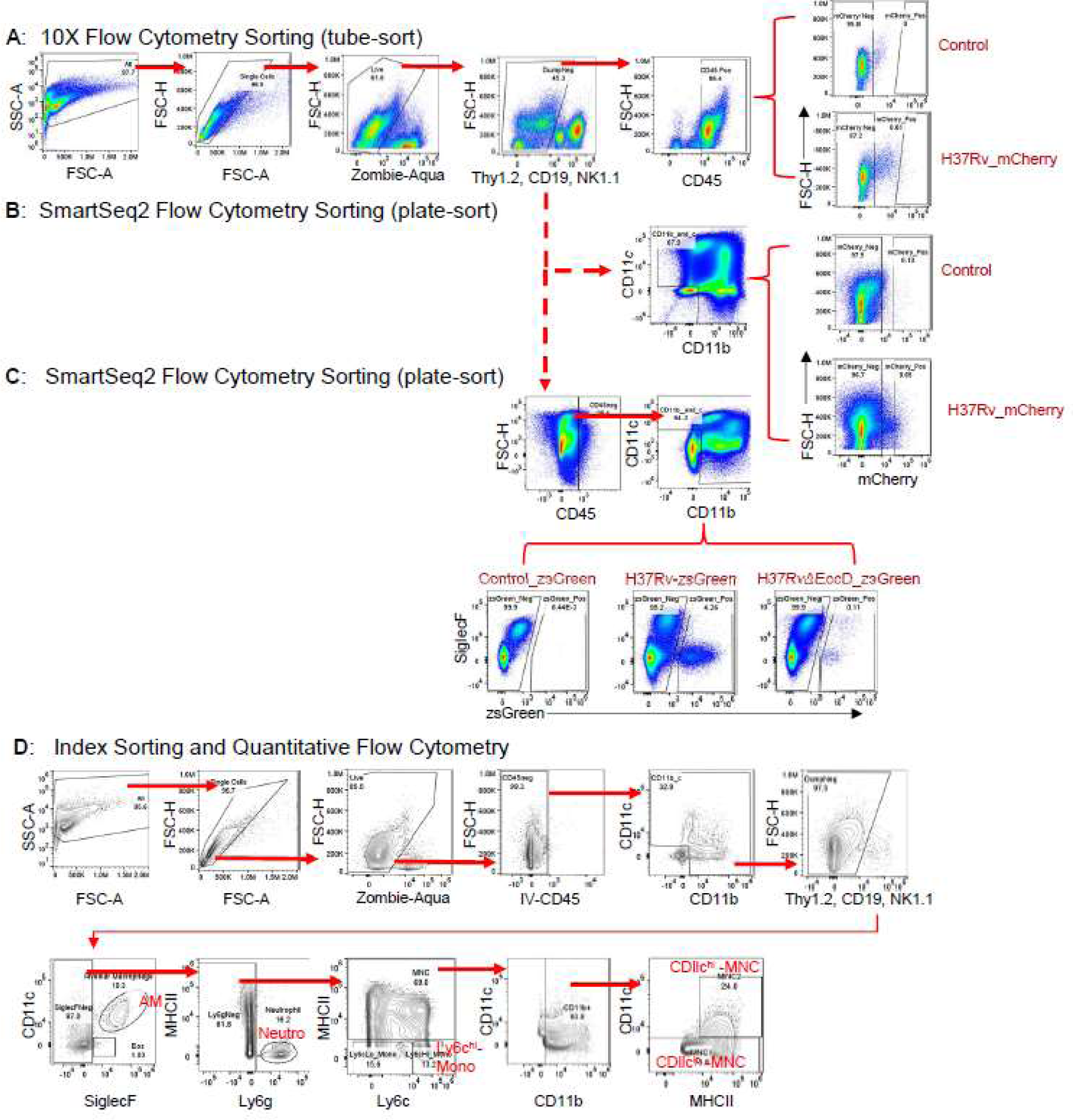
Flow cytometry sorting scheme to obtain lung mononuclear phagocyte transcriptional libraries in mice infected with MTB. **A)** Prior to 10X Chromium loading, live single cells were sorted on CD45 positivity, removing NK, T and B cells, into separate tubes based on mCherry positivity. **B)** Cells were sorted on CD11c and/or CD11b positivity for a more narrowed analysis, with mCherry positive or negative sorted separately into 384 well plates. **C)** In the setting of H37RvΔEccD infection, zsGreen fluorophore was utilized for improved demarcations. Intravascular monocytes were further assured removal through the addition of intravascular CD45. **D)** Schematic for quantitative flow cytometry, as well as stains for index sorting mean fluorescent intensity analysis used in panels B and C. Cells were subset into alveolar macrophages (AM; CD11b^lo^CD11c^hi^SiglecF^hi^), neutrophil (neutro; SiglecF^-^Ly6G^hi^CD11b^hi^), CD11c^lo^-mononuclear cell (MNC; CD11b^+^CD11c^lo^MHCII^+^) and CD11c^hi^-mononuclear cell (MNC2; CD11b^+^CD11c^hi^MHCII^hi^). Further analysis based on MTB-fluorescent protein expression was also completed. For all experiments, lung perfusion was performed prior to processing.

**Supplemental Figure 2:**
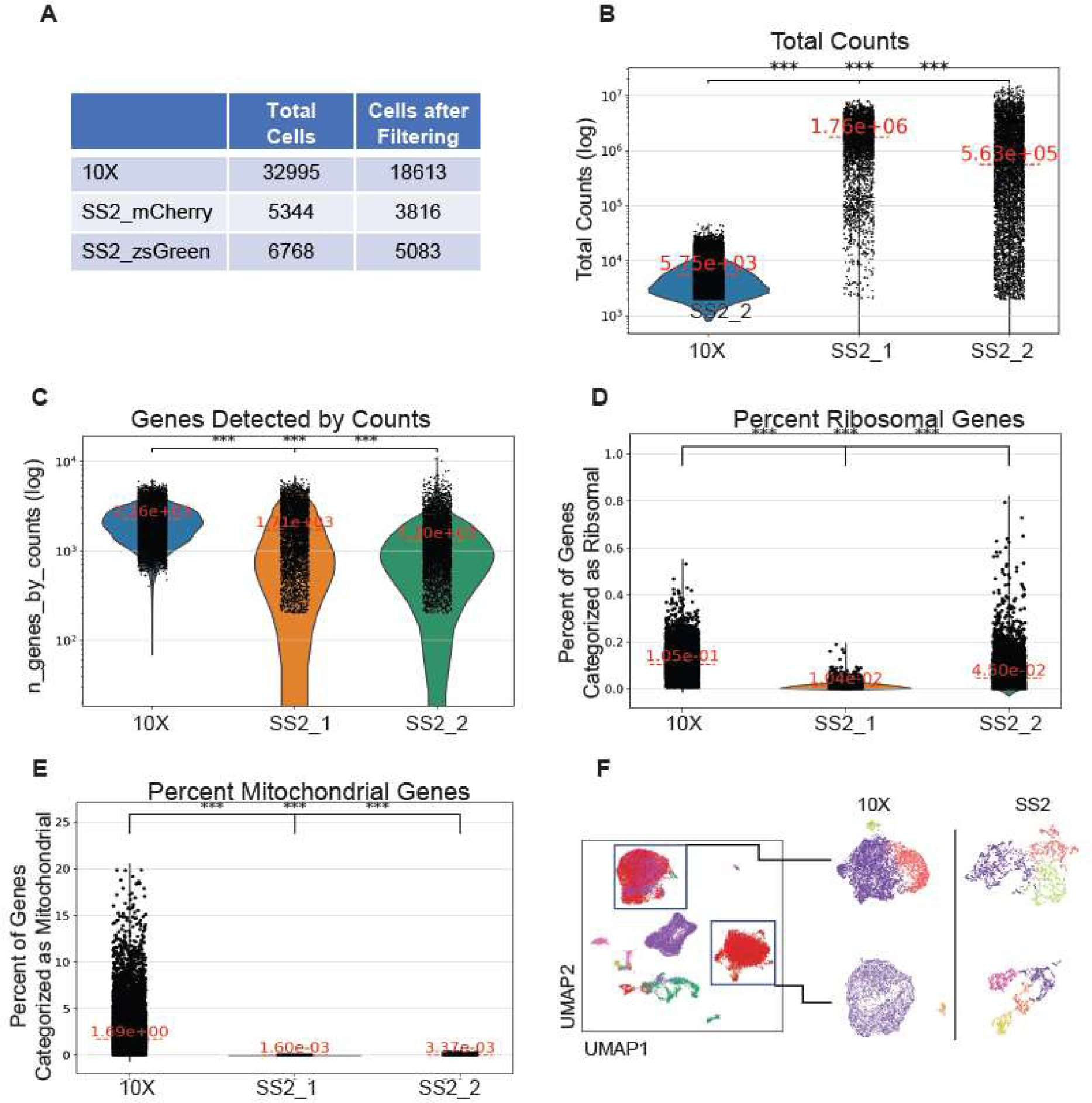
SmartSeq2 provides improved library depth when analyzing lung mononuclear phagocytes in mice infected with MTB within a BSL3 facility. **A**) Cell counts from the 3 single cell experiments. **B**) Total counts, **C**) genes (as normalized by counts), and **D**) ribosomal counts compared to each condition, where ***p<0.001. **F)** Using cellxgene, the two macrophage clusters were extracted by experimental method (i.e. 10X or smartseq2) and clustered using Leiden at a resolution of 0.2 (differential color delineation is per subcluster).

**Supplemental Figure 3:**
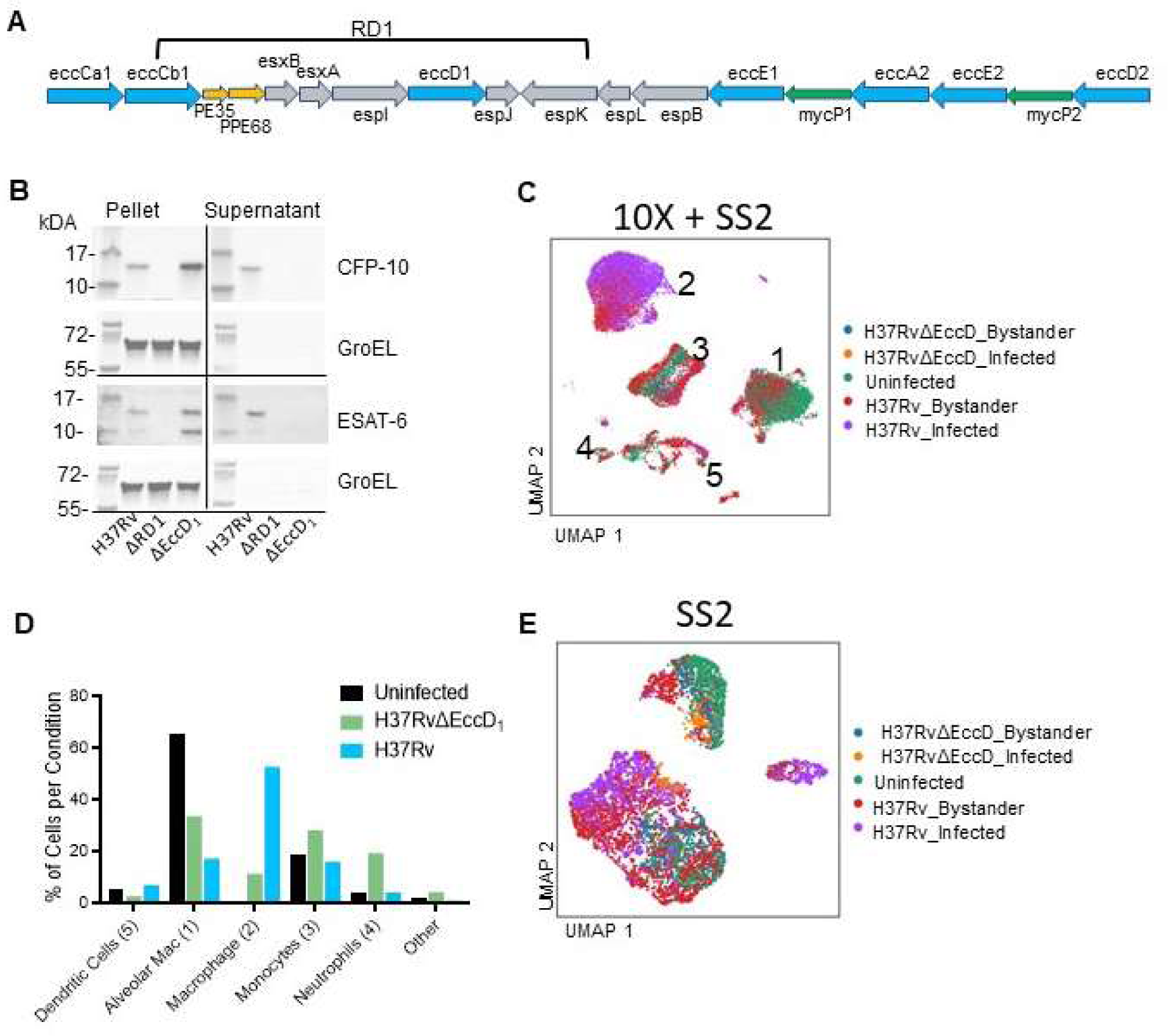
Macrophages are recruited differentially with or without ESX-1 presence. **A**) ESX-1 is a multi-protein complex encoded throughout, as well as outside, the RD1 operon. The RD1 operon also encompasses other genes not related to ESX-1. **B)** Representative immunoblots of the pellet and supernatant of MTB strains grown in minimal media was used to evaluate secretion of ESX-1 dependent CFP-10 and ESAT-6 in H37Rv, H37RvΔRD1, and H37RvΔEccD_1_ strains. **C)** Dimensionally reduced integrated data from 3 separate single-cell experiments depicts infected states, where ‘infected’ was determined by MTB-fluorescent probe expression on flow sorting, ‘bystander’ where a cell obtained from an MTB-infected mouse had no probe expression, and ‘uninfected’ cells from mice not exposed to MTB. **D)** Quantification of cell types recruited per condition, with percentage representative of that condition (ex. % macrophages over all cells in H37Rv-infected mice). Macrophage cell types are differentiated from alveolar macrophages, recruited macrophages, and monocytes as labeled in panel C. **E)** Removal of 10X dataset with reclustering of cells obtained from SmartSeq2 (Scanpy).

**Supplemental Table 1:** Final cell numbers and proportions. After quality control, cell-type identification, and subsetting from the larger atlas, cells obtained for SmartSeq2 were analyzed for proportion in each condition. Statistical analysis was conducted using a 2-sample chi-square with continuity correction).

| **Cell Type** | **Uninfected**<br>**(% Total)** | **H37RvΔEccD<sub>1</sub>**<br>**(% Total)** | **H37Rv**<br>**(% Total)** | **H37Rv versus**<br>**H37RvΔEccD<sub>1</sub> (p-**<br>**Val)** |
| --- | --- | --- | --- | --- |
| **moDerived**<br>**Macrophage** | 0<br>(0) | 83<br>(11.6) | 1656<br>(54.01) | <0.0001 |
| **Transitional**<br>**Monocyte** | 524<br>(43.49) | 289<br>(40.45) | 683<br>(22.28) | <0.0001 |
| **Alveolar**<br>**Macrophage** | 610<br>(50.62) | 297<br>(41.6) | 295<br>(9.62) | NS |
| **Notch<sup>Hi</sup>** | 0<br>(0) | 6<br>(0.84) | 273<br>(8.9) | <0.0001 |
| **Dendritic Cell** | 71<br>(5.89) | 39<br>(5.46) | 159<br>(5.19) | NS |

**Supplemental Figure 4:**
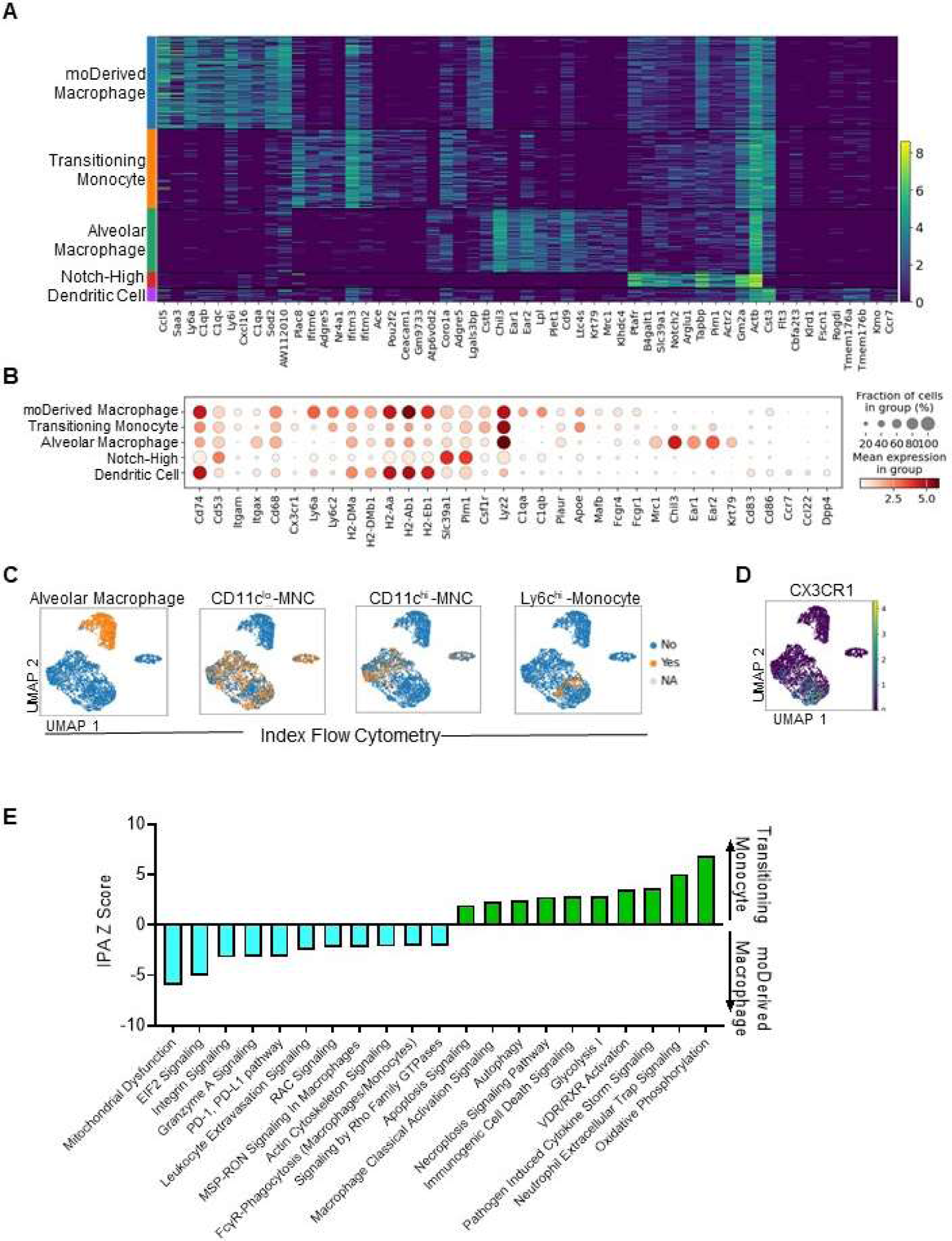
MTB induces recruitment of two large monocyte-derived cell subsets and one small mature dendritic cell cluster. C57BL/6 mice were infected with H37Rv or H37RvΔEccD_1_ for 28 days and single cell suspensions were sorted and processed using SmartSeq2 (Figure 2). **A)** Heatmap of 10 highest differentially expressed genes between clusters. **B)** Transcriptional canonical markers confirming subset delineation naming. **C)** Index sorting obtained during the acquisition of single cells into 384 well plates. **D)** Cx3cr1 gene expression of individual cells, demonstrating highest expression in the immature monocyte derived subset. **E)** Differential gene expression was conducted by MAST to compare recruited transitioning monocyte and monocyte-derived macrophages. Those with FDR <0.05 were used for Ingenuity Pathway Analysis. Shown are pathways with FDR <0.05 removing redundancy.

**Supplemental Table 2:**
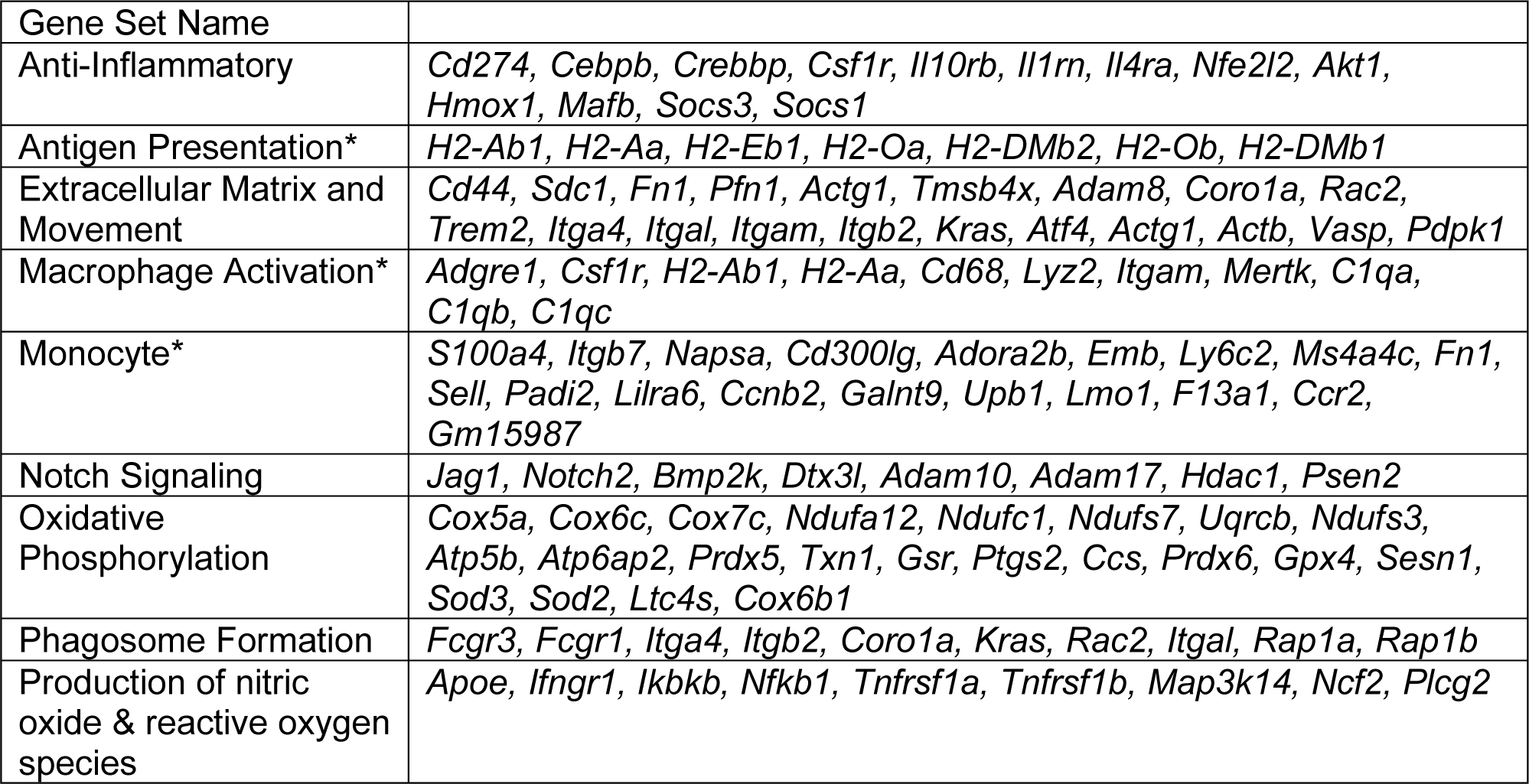
Genes utilized for each score name (Figure 3). *Denotes scores that were exhibited in *Sanin et al*.^54^.

**Supplementary Figure5:**
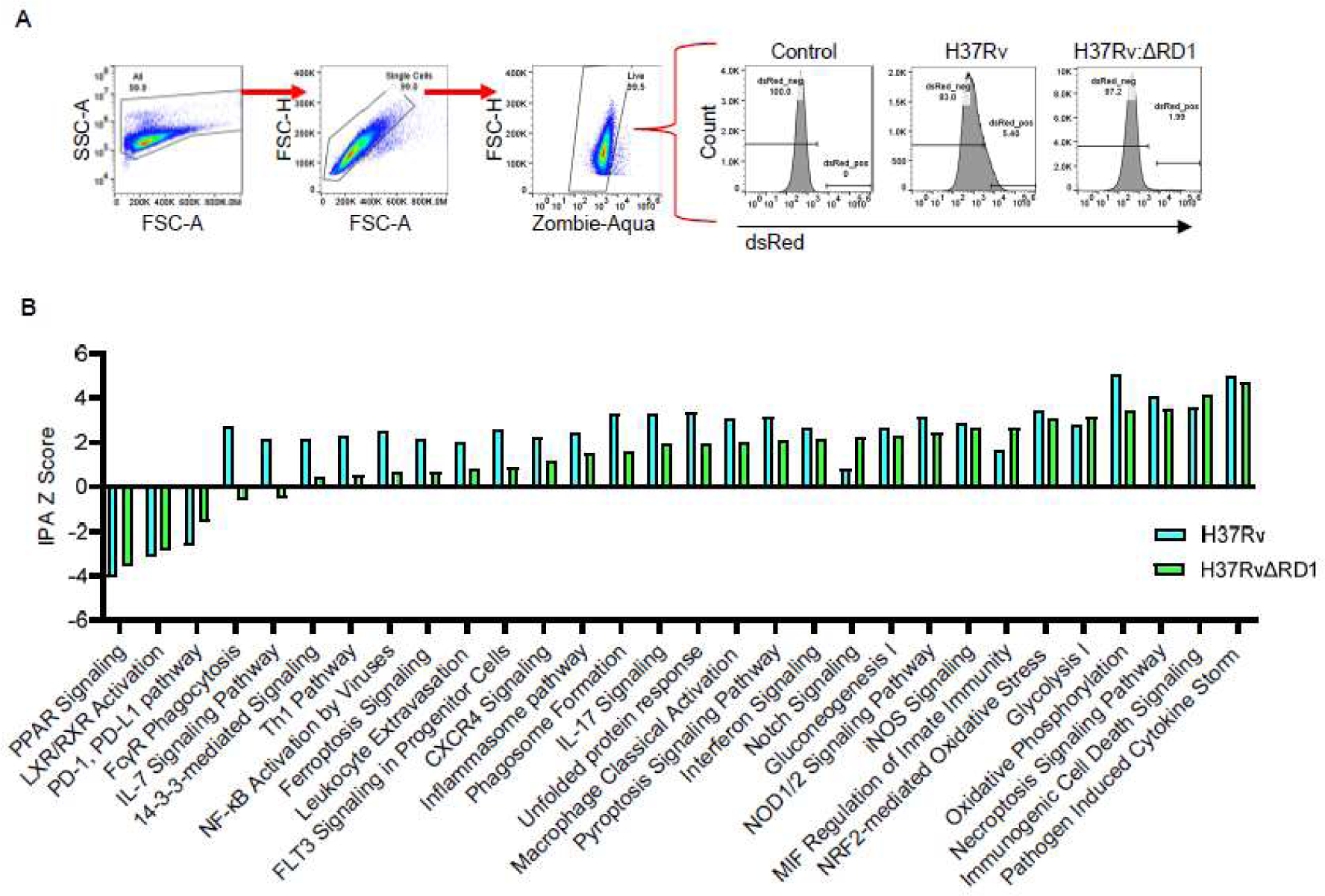
**A)** Flow cytometry sorting scheme to infected bone-marrow derived macrophages. Cells were infected with H37Rv or H37RvΔRD1 expressing dsRed for 24 hours. Cells were collected, washed, and stained for live/dead delineation using zombie-aqua, filtered into a single cell suspension, and sorted in a BSL3 contained Sony MA900. Shown are representative images to obtain single live cells, with cutoffs for dsRed positivity to delineate infected from uninfected cells. **B)** Qiagen Ingenuity pathway analysis was queried using differentially expressed genes determined from MAST analysis of infected cells compared to control cells, with an FDR <0.05. Shown are significantly enriched pathways scaled on Z score for directionality analysis, with >|2| considered significant.

**Supplemental Figure 6:**
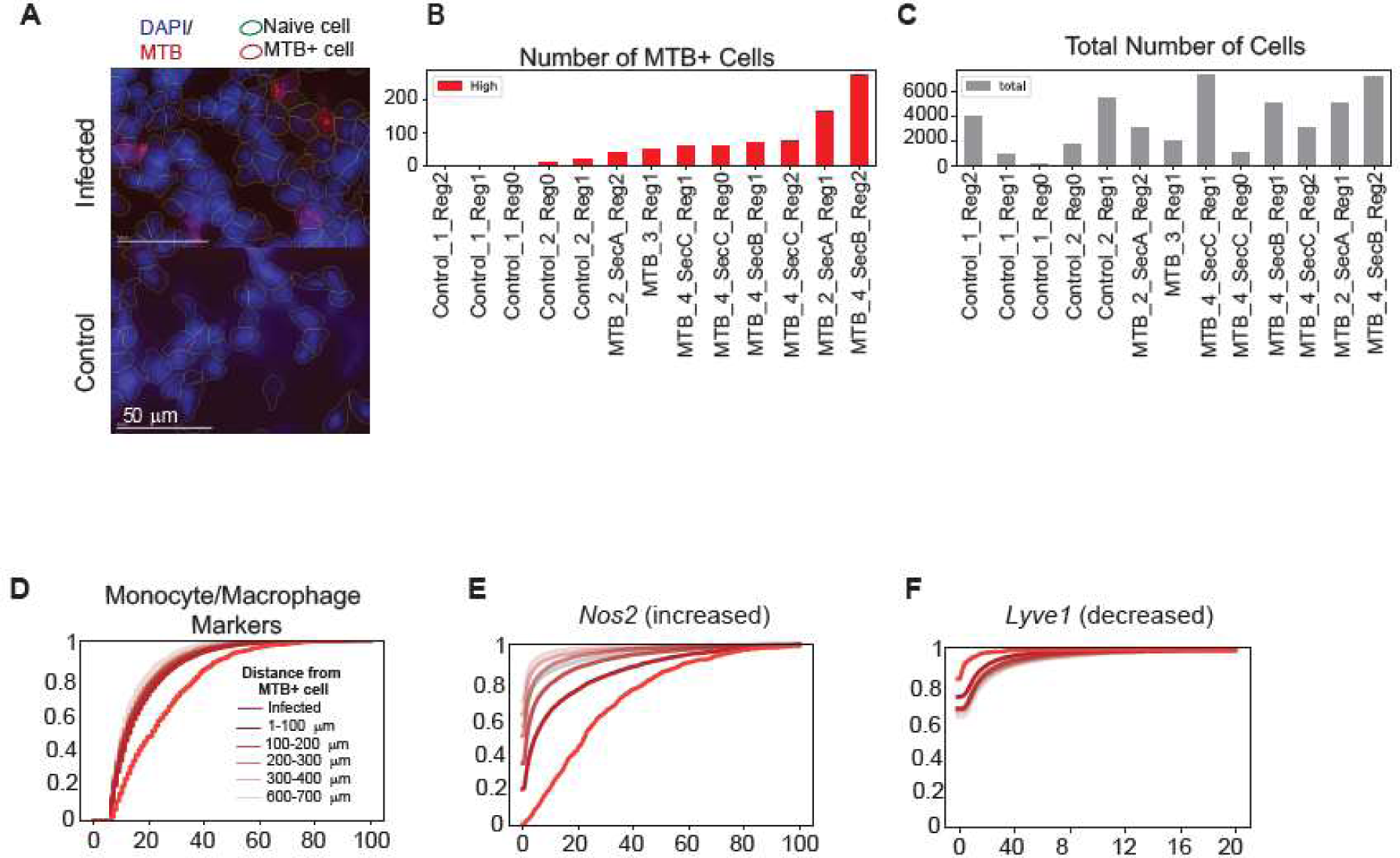
**A)** Identification of MTB positively infected cells in tissue slices obtained from control (uninfected) and H37Rv-infected C57BL/6 mice. MTB infected cells were identified by averaging the brightest 100 pixels from Cellbound3 (MTB antibody) channel per cell. Any cell in which this average was 300 or greater with a standard deviation in the MTB antibody channel greater than 150 was labelled as infected. Cells with lower standard deviations represented bright circular artifact staining and were not counted. Any cells that were outliers in space (lacking 2 or more infected neighbors within half a millimeter) or in brightness (>5 million average cell bound 3 intensity) were also removed. Regions with obvious tissue defects that produced large artifacts in the cellbound3 channel (i.e. tears) were excluded. **B)** Number of MTB positively infected cells in tissue slices, with **C)** total cell counts per section for comparison. **D**) Cumulative distribution plot of strictly defined monocyte/macrophages (*C1qb, C1qc, Clec4a2, Dnase1l3, Ms4a7, Sell, Trem2, Mmp9, Clec43*). In addition, cumulative distribution plots for **E)** *Nos2* and **F)** *Lyve1* within strictly defined monocyte/macrophages showing overall concordance with Fig6.

